# Diabetes alters neuroeconomically dissociable forms of mental accounting

**DOI:** 10.1101/2024.01.04.574210

**Authors:** Chinonso A. Nwakama, Romain Durand-de Cuttoli, Zainab M. Oketokoun, Samantha O. Brown, Jillian E. Haller, Adriana Méndez, Mohammad Jodeiri Farshbaf, Y. Zoe Cho, Sanjana Ahmed, Sophia Leng, Jessica L. Ables, Brian M. Sweis

**Affiliations:** Nash Family Department of Neuroscience, Friedman Brain Institute, Icahn School of Medicine at Mount Sinai, New York, NY 10029; Medical Scientist Training Program, Icahn School of Medicine at Mount Sinai, New York, NY 10029; Department of Biology, The University of Scranton College of Arts and Sciences, Scranton, PA 18510; Department of Chemistry, Barnard College of Columbia University, New York, NY 10027; Macaulay Honors at CUNY Hunter, New York, NY 10023; Hunter College High School, New York, NY 10128; Department of Psychiatry, Icahn School of Medicine at Mount Sinai, New York, NY 10029

## Abstract

Those with diabetes mellitus are at high-risk of developing psychiatric disorders, yet the link between hyperglycemia and alterations in motivated behavior has not been explored in detail. We characterized value-based decision-making behavior of a streptozocin-induced diabetic mouse model on a naturalistic neuroeconomic foraging paradigm called Restaurant Row. Mice made self-paced choices while on a limited time-budget accepting or rejecting reward offers as a function of cost (delays cued by tone-pitch) and subjective value (flavors), tested daily in a closed-economy system across months. We found streptozocin-treated mice disproportionately undervalued less-preferred flavors and inverted their meal-consumption patterns shifted toward a more costly strategy that overprioritized high-value rewards. We discovered these foraging behaviors were driven by impairments in multiple decision-making systems, including the ability to deliberate when engaged in conflict and cache the value of the passage of time in the form of sunk costs. Surprisingly, diabetes-induced changes in behavior depended not only on the type of choice being made but also the salience of reward-scarcity in the environment. These findings suggest complex relationships between glycemic regulation and dissociable valuation algorithms underlying unique cognitive heuristics and sensitivity to opportunity costs can disrupt fundamentally distinct computational processes and could give rise to psychiatric vulnerabilities.

---

Diabetes Mellitus, known colloquially as diabetes, refers to a group of metabolic diseases that impact an individual’s ability to regulate blood glucose via perturbations in insulin function, either insulin deficiency as in Type 1 diabetes, or impaired signaling through the insulin receptor as in Type 2 diabetes^1^. The brain is the body’s largest user of glucose as an energy source yet has often been considered as being spared from the effects of diabetes, despite literature to refute this^2^. Numerous studies have demonstrated that diabetes, even in the absence of signs of diabetes-associated vascular dysfunction, leads to altered cognitive performance and is associated with double the risk of psychiatric disorders as well as with neurodegenerative illnesses such as Alzheimer’s disease^3-9^. Postmortem brain tissue analysis of people with diabetes points toward alterations in key regions of reward circuitry that have been implicated in psychiatric sequelae, most notably the caudate, hippocampus, nucleus accumbens, and amygdala, as being disproportionately affected compared to other parts of the brain^10-15^. In animals, diabetes can augment central and peripheral catecholaminergic systems^16,17^. These findings suggest that reward processing might be disturbed, providing a possible link to increased psychiatric disorders among people with diabetes. However, our understanding of how brain function that may be altered in diabetes gives rise to vulnerabilities in motivated behavior remains limited.

Reward-seeking behavior derives from multiple decision-making systems in the brain^18^. Although the aforementioned brain regions affected by diabetes play coordinated roles in overall limbic function, each circuit can contribute to separable valuation algorithms via fundamentally distinct computations that can uniquely go awry in different psychiatric disorders. Neuroeconomics is an emerging field of decision science that leverages complex approaches in behavior to quantify how the physical limits of the brain constrain the way cognitive mechanisms process reward-related information^19^. This encompasses characterizing multifactorial aspects of motivation that operationalize reward value along several dimensions, integrate choice processes with environmental circumstances, and capture brain-body interactions with evolutionarily conserved cognitive heuristics that may depend on, for instance, energy balance and metabolic demand^20-22^. Such approaches would be critically informative if applied toward translational studies of diabetes. Recent insights from neuroeconomic principles offer novel approaches to investigate decision-making information processing capable of resolving behavior into discretely measurable computational units in a manner that is biologically tractable and readily translatable across species^23-25^.

Here, we set out to examine how diabetes alters multiple aspects of value-based decision-making behavior. We characterized a rich dataset of neuroeconomic choice processes in streptozocin (STZ)-induced diabetic mice^26^ tested across a longitudinal naturalistic foraging paradigm, Restaurant Row^27,28^. In this complex behavioral task, mice must forage for their primary source of food while on limited daily time budget in a closed-economy system by navigating a maze with four uniquely flavored and contextualized feeding sites, or “restaurants.” Mice learn to associate the pitch of a tone with reward cost in the form of a delay required to obtain food. Mice are free to choose in a self-paced manner how to invest time in competing actions, whether to accept and wait for presented offers or skip and proceed to the next restaurant. Because time is a limited commodity, choices on this task can have dire opportunity costs and are interdependent both between trials and across days. Importantly, we experimentally manipulated not only the distribution of reward costs presented to the animals across weeks to months but also systematically varied the rate of the changing economic landscape of the environment. We discovered that diabetic mice employed altered behavioral strategies that not only develop across an increasingly reward-scarce environment but also depend on and integrate the history of its salience in order to drive changes in fundamentally distinct valuation algorithms. This work demonstrates the utility of applying neuroeconomic approaches to animal behavior to uncover dissociable motivational processes capable of altering decision-making behavior in diabetes.

## Results

### STZ treatment induces chronic hyperglycemia as a mouse model of diabetes

In order to generate a mouse model of diabetes, 40 C57BL/6J male mice were treated with intraperitoneal injections for 5 consecutive days of either Hank’s buffered saline solution (HBSS) as the vehicle (VEH) control group or low-dose streptozotocin (STZ, 50 mg/kg), which ablates insulin-producing beta cells of the pancreas^26^ (Fig. 1a). Body weight and morning fasting blood glucose levels sampled from the tail vein were monitored weekly over the next 7 weeks to allow for the effects of chronic hyperglycemia to incubate while mice remained group-housed with ad libitum access to regular chow. We found that treatment with STZ induced a robust and long-lasting increase in blood glucose levels accompanied by a transient decrease in body weight that briefly delayed subsequent weight gain compared to VEH-treated mice (Fig. 1b-c). Prior to starting the behavioral task, blood HbA1c was measured to confirm diabetic phenotype. We found STZ-treated mice displayed significantly elevated HbA1c levels, confirming that these animals were chronically hyperglycemic compared to VEH-treated mice (Fig. 1d). Next, mice were single-housed and food-restricted to between 80-85% of free feeding weight as they began longitudinal testing on the Restaurant Row paradigm.

**Fig. 1.**
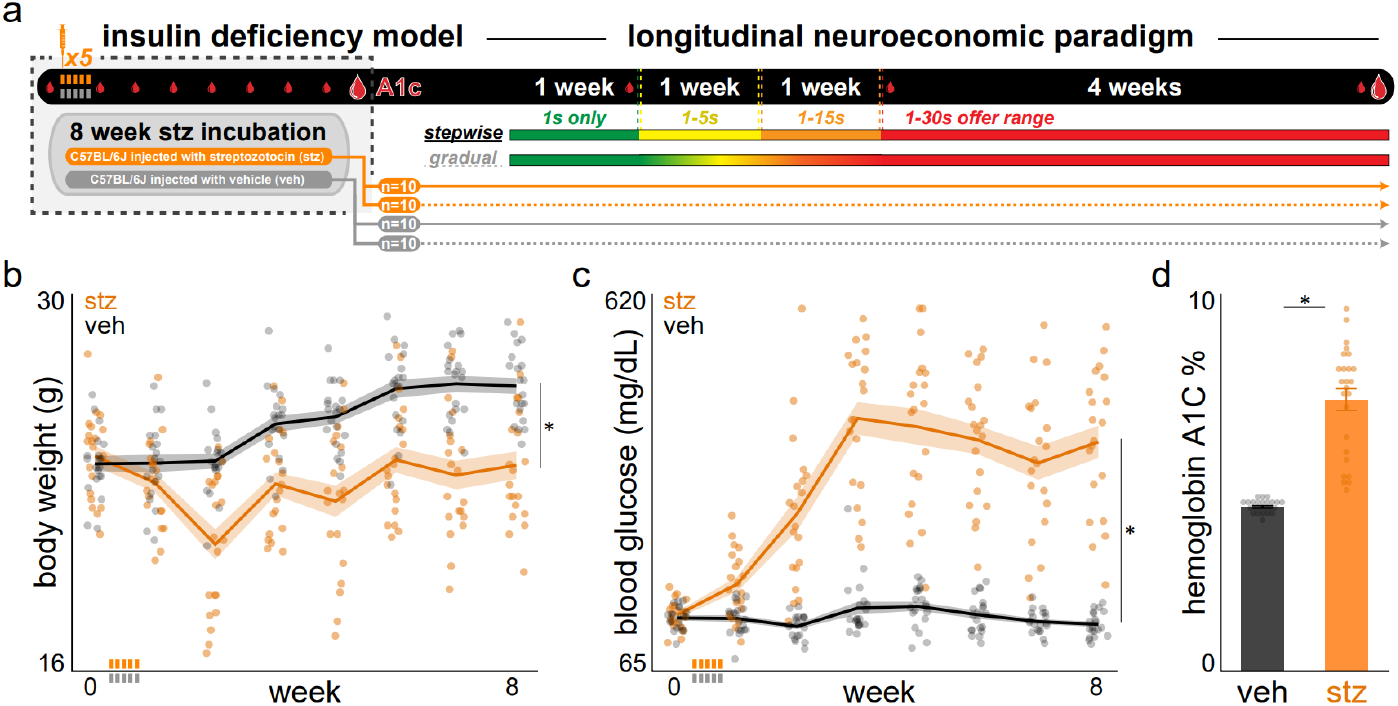
Streptozocin-induced diabetes mellitus mouse model of chronic hyperglycemia. **a.** Experimental timeline. Dashed box indicates time period relevant for this figure: hyperglycemia incubation. Randomly selected male C57BL/6J mice received daily intraperitoneal injections of vehicle (veh, Hank’s balanced saline solution, n=20) or streptozocin (stz, 50 mg/kg, n=20) injections for 5 consecutive days (gray or orange tick icon). STZ is an antibiotic that ablates insulin-producing beta cells of the pancreas. Mice were allowed to incubate for 8 weeks while sampling body weights and fasting tail vein blood glucose levels (small droplet icon) weekly before sampling tail vein blood Hemoglobin A1c once (large droplet icon) and then beginning longitudinal neuroeconomic behavioral testing for an additional ∼2 months. **b** Weekly body weights. As expected, STZ-treated mice experienced an initial drop in weight before returning to rate of weight gain similar to VEH mice (*F*=19.237, *p*<0.0001). **c** Fasting tail vein blood glucose levels. STZ treatment significantly elevated blood glucose measurements compared to VEH-treated mice (*F*=73.841, *p*<0.0001). **d** Tail vein blood Hemoglobin A1c obtained at week 8, confirming chronic hyperglycemia in STZ-treated mice, sampled 3 days before starting behavioral testing (*t*=9.96, *p*<0.0001). *Represents significant differences between veh / stz groups. Dots represent individual mice. Shading / error bars represent ±1 SEM.

### STZ-treated mice are able to acquire the basic structure of the Restaurant Row task

During the first week of behavioral testing on the Restaurant Row task, all reward offers cost only 1 s (Fig. 2a-b). During this period, all mice quickly acquired directionality of the task with no differences in number of laps run in the correct counter-clockwise direction between STZ- and VEH-treated mice (Fig. 2c). We also found no differences in travel time when running in the hallways between restaurants among STZ- and VEH-treated mice (Fig. 2e). Despite acquiring running in the correct direction at the same rate and stabilizing at similar number of laps, we found a modest decrease in the total number of pellets earned in STZ-treated mice compared to VEH-treated mice at the end of the first week of behavioral testing (Fig. 2d). This effect negatively correlated with HbA1c levels but did not correlate with body weight, reflecting a change in foraging behavior that is likely mediated by their hyperglycemic profile (Supplementary Fig. 1).

**Fig. 2.**
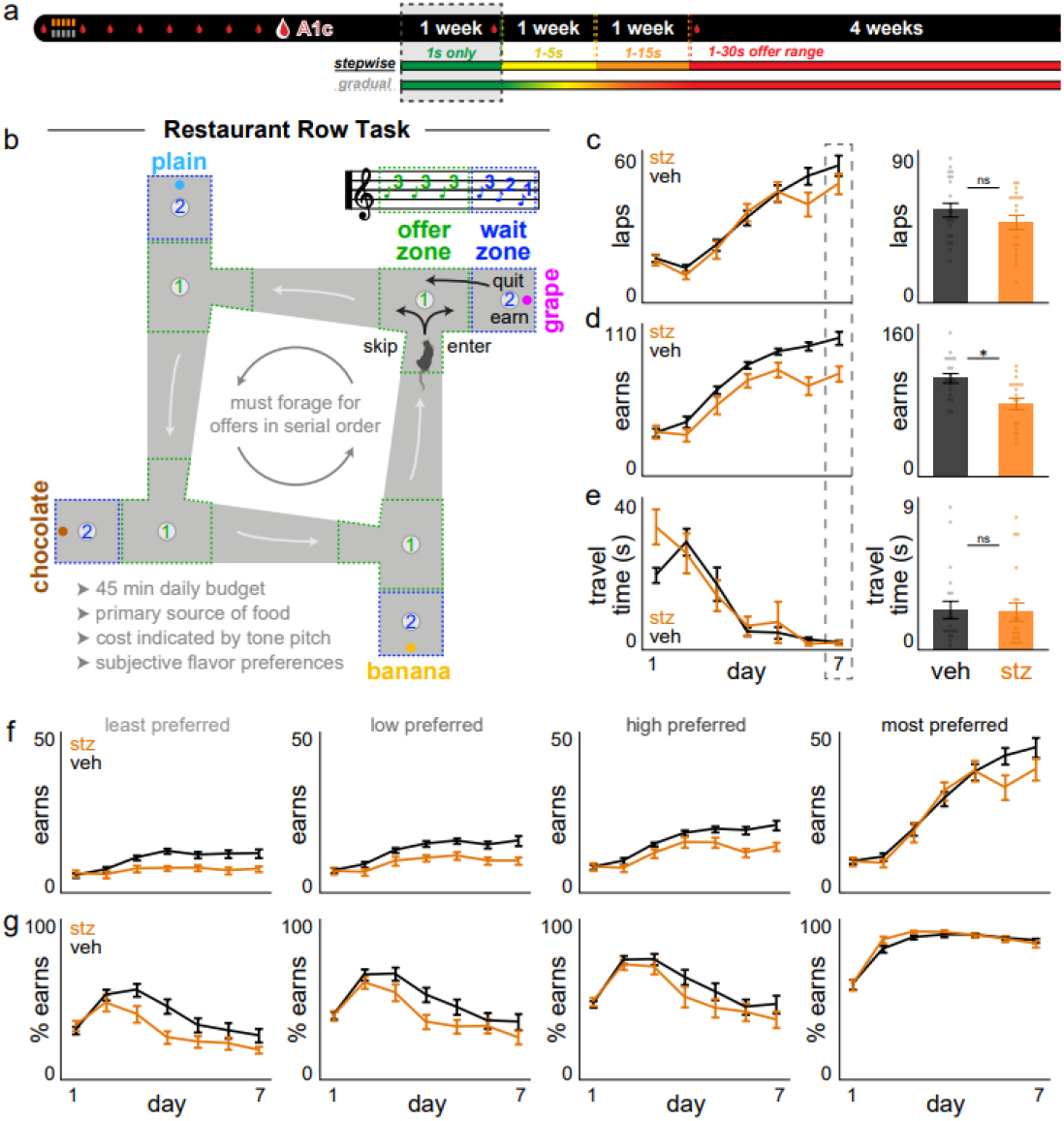
STZ-treated mice are able to acquire the basic structure of the naturalistic, neuroeconomic foraging task Restaurant Row. **a** Timeline. Dashed box indicates time period relevant for this figure: first week of behavioral testing. **b** Task schematic. Mice were allotted 45 min daily to invest time foraging for their primary source of food in a self-paced manner. Costs to obtain rewards were in the form of delays mice would have to wait near feeder sites. Mice were required to run in a counterclockwise direction encountering offers for different flavors at each “restaurant” in serial order. Each restaurant, separated by hallways, was divided into a T-shaped “offer zone” choice point and a separate “wait zone” that housed the pellet dispenser. Upon offer zone entry from the correct heading direction, a tone sounded whose pitch indicated the delay mice would have to wait if accepting the offer by entering the wait zone. If entered, tone pitch descended in the wait zone, cuing the indicated delay. Each trial terminated if mice skipped in the offer zone, quit during the countdown in the wait zone, or earned a reward, after which animals were required to proceed to the next restaurant. **c-g** Simple behavioral metrics across the first week of testing during which all offers were 1 s only (lowest pitch, 4 kHz): (c) laps run in the correct direction, (d) total rewards earned, (e) inter-trial travel time between restaurants, (f) earnings split by flavors ranked from least to most preferred by summing each day’s end-of-session totals in each restaurant and (g) normalized to number of laps run (treatment x rank: *F*=3.823, *p*<0.01). In (c-e) right, *represents significant differences between veh / stz groups on day 7 (dashed box): laps: *t*=1.24, *p*=0.223; earns: *t*=3.53, *p*<0.01, travel time: *t*=0.17, *p*=0.863. Dots represent individual mice. Error bars represent ±1 SEM.

### STZ-treated mice display altered flavor-specific meal consumption patterns

Upon closer examination of rewards earned during the first week, we found that differences in food intake between STZ- and VEH-treated mice were largely driven by disproportionately fewer earnings for less preferred flavors, stabilizing by day 7 (Fig. 2g). These data indicate STZ-treated mice exhibited changes in revealed preferences that were asymmetrically skewed rather than globally shifted among flavors. To characterize meal consumption patterns, we next calculated earnings across the 45 min session segregated into 2.5 min bins (Fig. 3a). Overall, we found that both groups earned fewer rewards across the session (Fig. 3b), reflecting a measure of within-session satiety. STZ-treated mice, however, displayed this decrease to a greater degree (Fig. 3c-d), despite earning amounts of food in the first 2.5 min bin similar to VEH-treated mice in each restaurant (Fig. 3c). When examining within-session meal patterns by restaurant, we found that decreased earnings across the session were largely driven by the most preferred flavor for both the VEH- and STZ-treated groups (Fig. 3c). That is, on day 7, satiety-related effects were only observable in the most preferred restaurant. Thus, earnings for less preferred flavors remained relatively flat within-session, albeit downshifted in STZ-treated mice (Fig. 3c). When calculating meal patterns normalized to total food earned, we found that STZ-treated mice consumed more of their day’s proportion of the most preferred flavor earlier in the session and prioritized over other flavors compared to VEH-treated mice (Fig. 3e). These data suggest that elements of satiety, which may be augmented by STZ treatment, interact with more intricate aspects of value-based decision-making and goal-directed behavior while foraging. To examine to what extent hyperglycemia might contribute to these effects, we measured body weight and blood glucose pre- and post-task. As expected, blood glucose increased post-task in both groups, with STZ-treated mice remaining hyperglycemic despite being food-restricted compared to VEH-treated mice (Fig. 3f-g). Change in blood glucose positively correlated with change in body weight in STZ-but not VEH-treated mice, consistent with dysregulation of glucose homeostasis in response to a meal (Fig. 3h). Pre-task blood glucose negatively correlated with earns for both least- and most-preferred flavors in STZ-treated mice on day 7 (Fig. 3i-j). Next, to investigate dissociable valuation processes that may be altered in these mice, we characterized how economic decision strategies develop as task complexity and environmental demand increases.

**Fig. 3.**
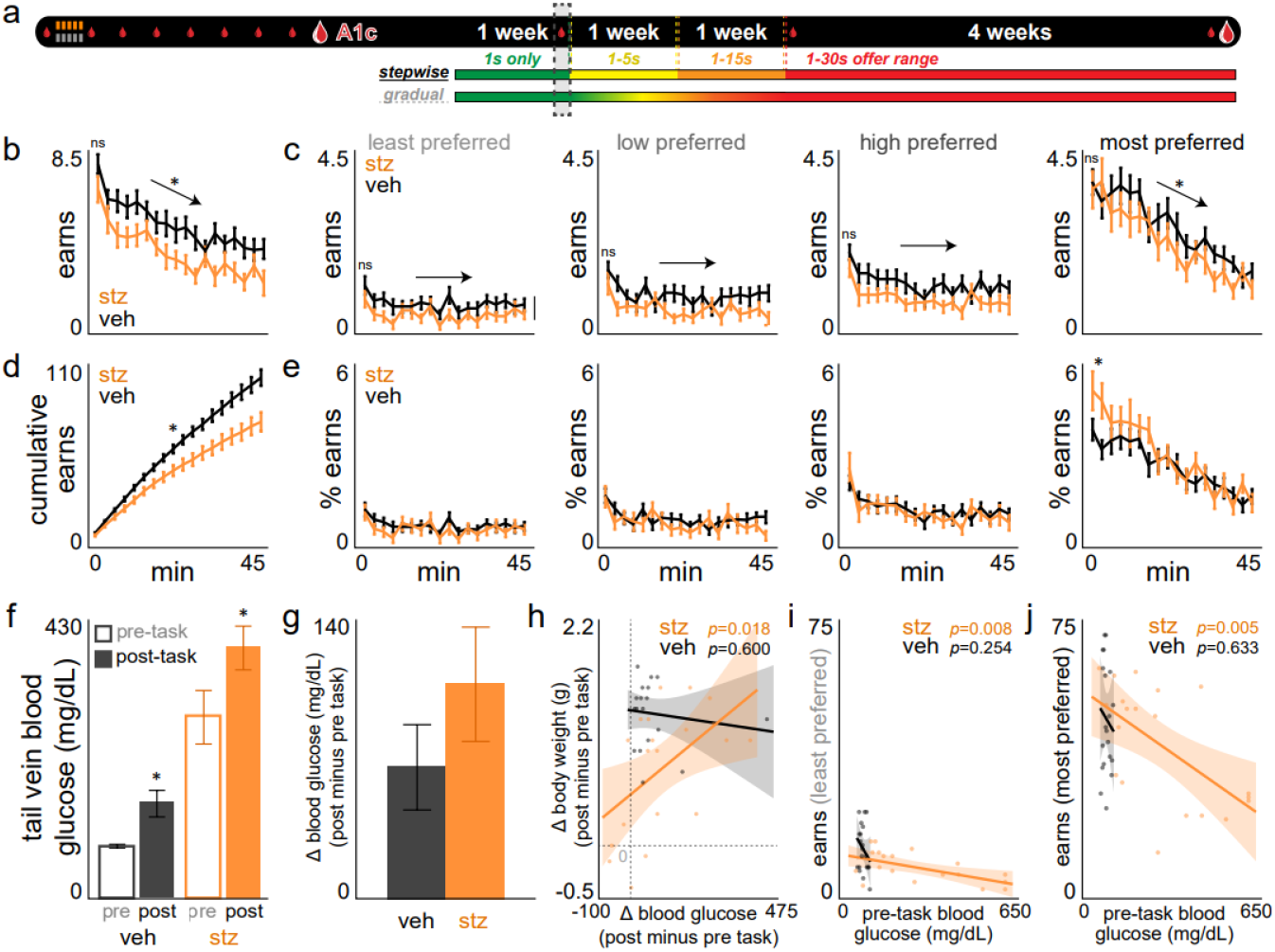
STZ-treated mice display early satiety and within-session meal consumption patterns skewed toward most preferred flavors. **a** Timeline. Dashed box indicates time period relevant for this figure: day 7. **b-c** Rewards earned within each 2.5 min bin across the session (b) in total earnings (main effect of session time on total earnings across groups: *F*=142.022, *p*<0.0001) or (c) earnings split by flavor ranking (no differences between groups in the first time bin: *F*=0.047, *p*=0.987; flavor x session time: *F*=74.690, *p*<0.0001). **d** Cumulative total rewards earned summed across the session (treatment x session time: *F*=12.469, *p*<0.01; midsession: *t*=3.13, *p*<0.01). **e** Percentage of total session rewards earned split by flavor (first time bin: treatment x rank: *F*=5.317, *p*<0.01). **f** Tail vein blood glucose levels sampled immediately before and after day 7’s session (main effect of time point pre vs. post task: *F*=23.915, *p*<0.0001; main effect of treatment: *F*=36.666, *p*<0.0001). **g** Change in blood glucose from (f) post minus pre task. **h** Scatter plot of change in blood glucose from (g) against change in body weight measured immediately before and after day 7’s session. Gray dashed lines indicate 0 on both axes. **i-j** Scatter plot of pre-task blood glucose from (f) against day 7’s end-of-session earns for (i) least and (j) most preferred flavors. *Represents significant differences between veh / stz groups. Dots represent individual mice. Error bars represent ±1 SEM. Shading represents 95% confidence interval of linear fit.

### Mice learn to forage in a stepwise or gradually increasingly reward-scarce environment

A feature of the Restaurant Row task is the longitudinal nature of its closed-economy system in which behavior is interdependent across days as animals work for their primary source of food. Increasing the distribution of offer costs while animals remain on a fixed, limited time budget can elicit an economic challenge^21^. Because such a challenge can be metabolically demanding particularly for mice that may have differing energetic needs such as these STZ-treated mice, we experimentally manipulated the rate of change of reward scarcity in the environment in two ways. After the first week of testing during which all offers remained at 1 s only (epoch 1: a relatively reward-rich environment), groups of mice were split and advanced to the next stage of testing where the range of offers available in the task environment increased via one of two schedules: (i) stepwise or (ii) gradual (Fig. 4a, Supplementary Fig. 2). These schedules elicited a relative decrease in earned rewards that was either stepwise or gradual, respectively, across days 8-22 similarly for both STZ- and VEH-treated mice (Fig. 4b-c). Both schedules yielded matched earnings and reinforcement rate across groups of mice at each stepwise transition point and were equivalent by day 22 and beyond (Fig. 4b,d). This allows us to examine, *ceteris paribus*, not only how mice adapt choices to different rates of changing environments between days 8-22 but also how the experience and salience of the prior environmental change might bolster different valuation strategies once in the same, final reward-scarce environment after day 22. When examining meal consumption in the 1-30 s reward scarce environment split by restaurant, we found that STZ-treated mice displayed inverted meal patterns across the session and prioritized consuming more of their favorite flavor sooner compared to VEH-treated mice regardless of schedule history (Fig. 4e, Supplementary Fig. 3i-p). However, when calculating reinforcement rate by restaurant, we found a significantly higher latency between earns in STZ-treated mice compared to VEH-treated mice only after experiencing the gradual but not stepwise schedule that predominately affected less preferred flavors (Fig. 4f). These data suggest environmental history interacts with valuation strategies uniquely altered by STZ.

**Fig. 4.**
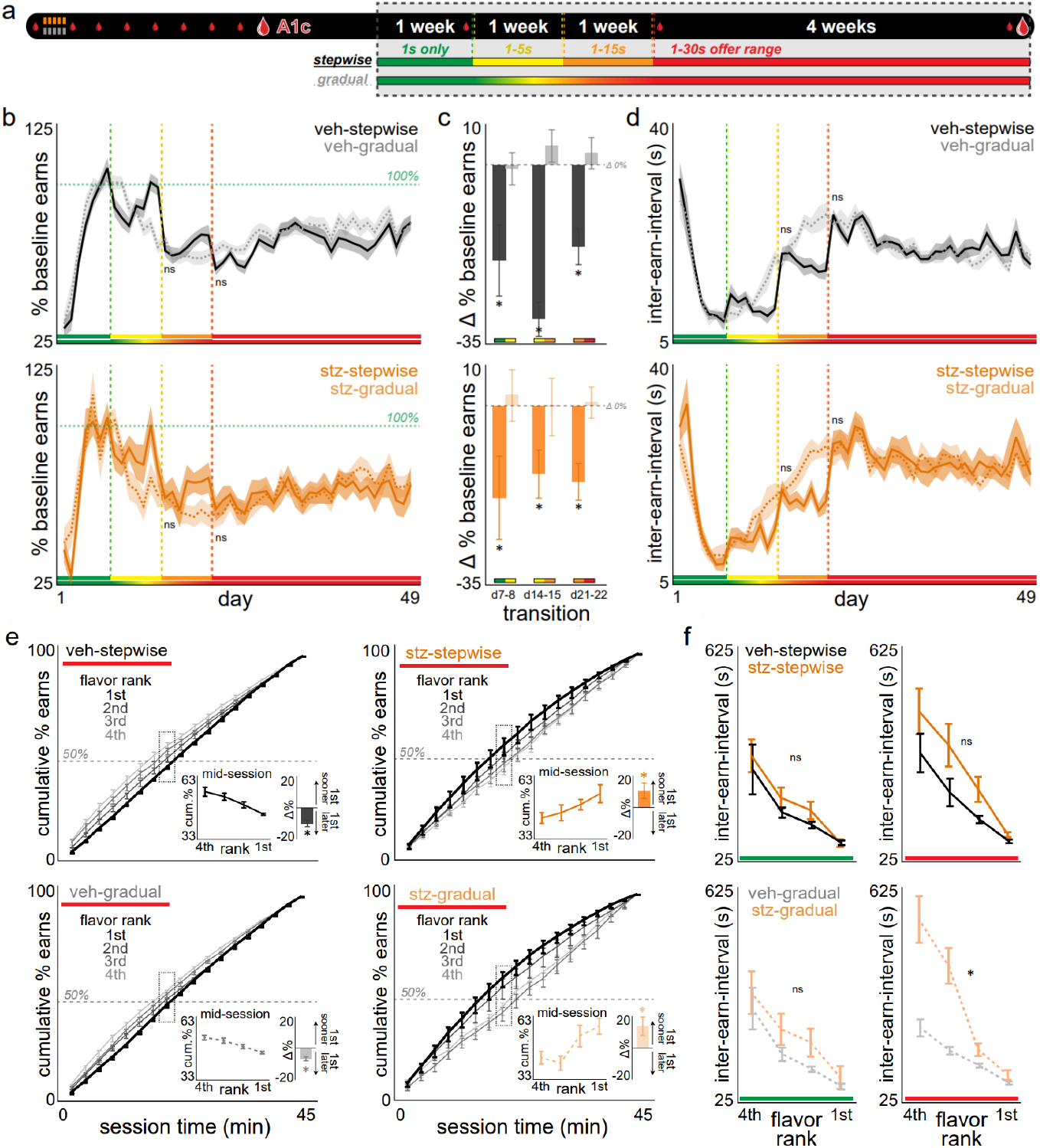
STZ-treated mice forage differently under budget constraints as reward availability decreases but depending on the salience of change in environmental scarcity. **a** Timeline. Dashed box indicates time period relevant for this figure: entire 7-week Restaurant Row paradigm. **b** Total rewards earned each day normalized to the average of days 5-7 earnings (termed 100%, horizontal dashed green line). Visual guidance color bars along the x-axis reflect the experimental schedules (stepwise or gradual) illustrated in the timeline in (a). Vertical dashed green-yellow-red lines indicate the transition points of the stepwise schedule (or matched days of the gradual schedule) and are re-used throughout all other figures as a visual aid. Top: VEH; bottom: STZ (treatment x schedule after transition points: *F*=0.937, *p*=0.335). **c** Change in (b) at each transition point of the stepwise schedule (or matched days of the gradual schedule): d7-8 (1 s only to 1-5 s offers, green to yellow), d14-15 (1-5 s to 1-15 s offers, yellow to orange), and d21-22 (1-15 s to 1-30 s offers, orange to red). Horizontal dashed gray line indicates 0 change (main effect of schedule: *F*=52.500, *p*<0.0001; no main effect of treatment: *F*=0.911, *p*=0.342). **d** Time elapsed between subsequent earns of any flavor (treatment x schedule after transition points: *F*=0.051, *p*=0.821). **e** Within-session cumulative earnings summed across 2.5 min bins normalized to end-of-session earnings for each restaurant. Data collapsed across the entire 1-30 s epoch (days 22-49, red). Horizontal dashed gray line indicates 50% of each flavor’s meal consumed. Dashed square box highlights mid-session data plotted within inset figure showing ranking spread (left) and a summary difference score (right) between most minus least preferred restaurants to depict whether favorite flavors are consumed relatively sooner (+) or later (-) in the session (main effect of treatment: *F*=25.326, *p*<0.0001). Horizontal black line indicates 0 difference. **f** Time elapsed between subsequent earns of the same flavor split by ranking. Data collapsed across either the 1 s only (green, treatment x rank: *F*=0.348, *p*=0.790) or 1-30 s epoch (red, treatment x rank: stepwise: *F*=0.752, *p*=0.525; gradual: *F*=9.779, *p*<0.0001). Shading / error bars represent ±1 SEM.

### STZ treatment alters economically distinct decision-making policies

To characterize economic decision patterns, next, we calculated the likelihood of selecting choices as a function of offer cost in both the offer zone (enter vs. skip) and wait zone (earn vs. quit) of each restaurant. We found that all mice were capable of discriminating tone pitch and could adhere to the basic economic structure of the Restaurant Row task, whereby interactions between choice outcomes and offer cost scaled with flavor preferences (Supplementary Fig. 3a-d). In order to approximate economic thresholds, or indifference points, of willingness to accept and earn rewards, we fit a heaviside-step regression to choice outcomes as a function of cost and measured curve inflection points each day across the entire experiment (Fig. 5a-d). This analysis revealed a large discrepancy in economic thresholds between offer zone and wait zone decision policies upon transitioning into the 1-30 s reward scarce environment. Overall, offer zone thresholds were significantly higher than wait zone thresholds, indicating mice were likely to accept offers more expensive than they were actually willing to wait for and earn. This discrepancy captures a metric of economic choice conflict between zones and was greater for more preferred flavors (Supplementary Fig. 4c). We discovered altered economic thresholds between STZ- and VEH-treated mice, but only in those tested on the gradual and not stepwise schedule and whose threshold direction of change depended on both the restaurant and zone type (Fig. 5b,d). Offer zone thresholds of less preferred flavors were lower in STZ-compared to VEH-treated mice (Fig. 5d). Wait zone thresholds of more preferred flavors were higher in STZ-compared to VEH-treated mice (Fig. 5b, see Supplementary Fig. 3e-h for examination of thresholds between days 8-22). These data indicate complex changes in distinct stages of the decision process within-trial - willingness to accept (offer zone) vs. willingness to wait (wait zone)- are differentially altered in STZ-treated mice.

**Fig. 5.**
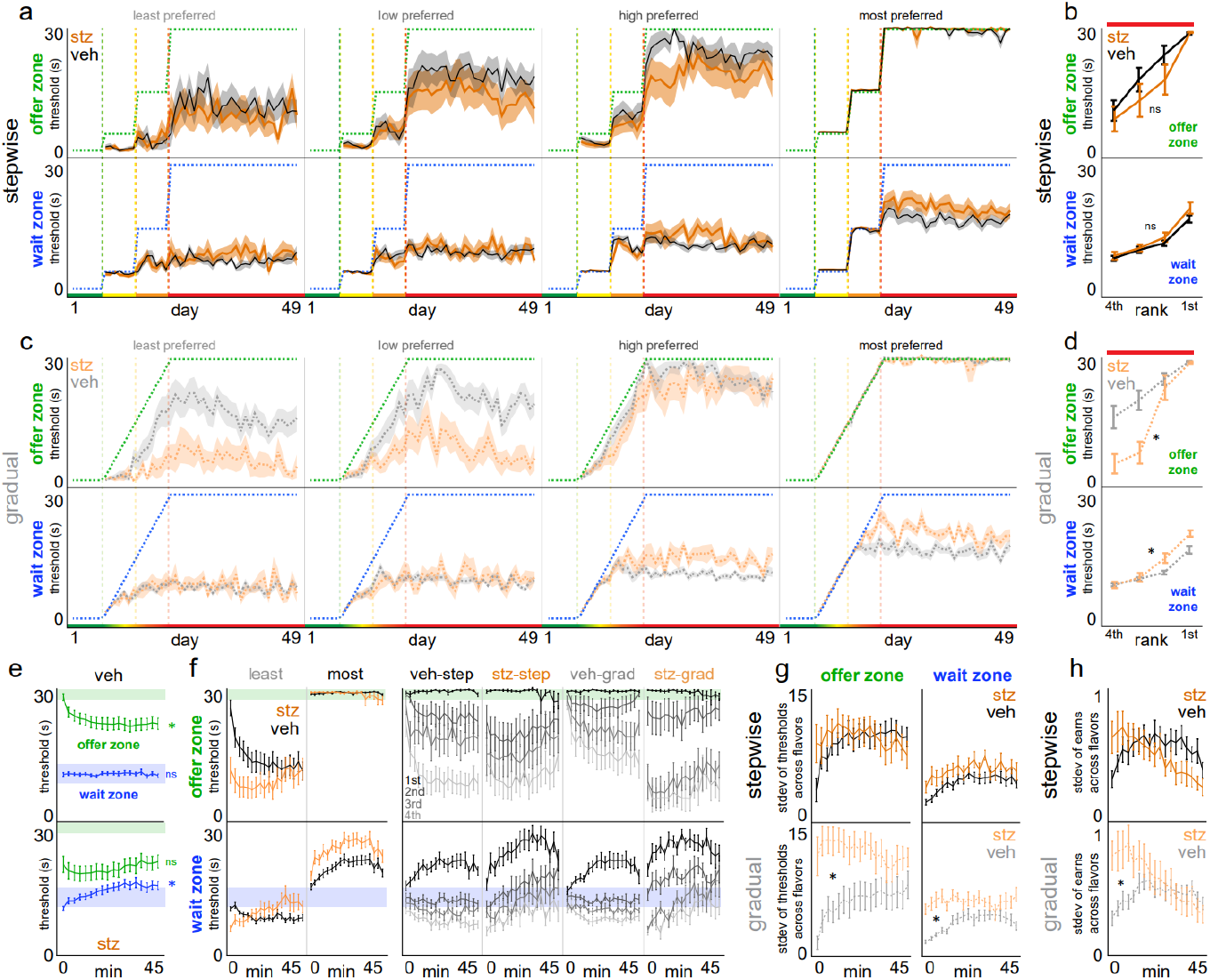
STZ-treated mice develop unique decision-making policies in fundamentally distinct types of choices. **a**-**d** Offer zone (top) and wait zone (bottom) thresholds plotted (a,c) each day across the entire Restaurant Row paradigm or (b,d) split by flavor ranking collapsed across the 1-30 s epoch. Green (offer zone) and blue (wait zone) dashed staircase represent the maximum possible threshold for stepwise (top) or gradual (bottom) schedules (offer zone>wait zone thresholds: *F*=74.715, *p*<0.0001; stepwise: offer zone treatment x rank: *F*=0.559, *p*=0.645; wait zone treatment x rank: *F*=0.572, *p*=0.635; gradual: offer zone treatment x rank: *F*=4.354, *p*<0.01; wait zone treatment x rank: *F*=3.390, *p*<0.05). **e-f** Offer zone and wait zone thresholds within each 2.5 min bin across the session (data collapsed from entire 1-30 s epoch). Both schedules collapsed in (e) depicting all VEH- (top) or STZ-treated mice (bottom). Horizontal shaded bands represent offer zone threshold of 30 (green) or wait zone threshold of 9.s-13.5 (blue), corresponding to thresholds representing a strategy that would yield the maximum amount of total food as one type of optimal strategy, should animals ignore flavors, that was previously theoretically and empirically determined^21^ (VEH: main effect of time on offer zone threshold: *F*=12.695, *p*<0.001, but not wait zone threshold: *F*=0.039, *p*=0.843; STZ: main effect of time on wait zone threshold: *F*=50.312, *p*<0.0001, but not offer zone threshold: *F*=3.577, *p*=0.060). Analysis from (e) replotted and split by least and most preferred flavors in (f) for both schedules collapsed (left, VEH: offer zone rank x time: *F*=7.241, *p*<0.0001, wait zone rank x time: *F*=22.096, *p*<0.0001; STZ: offer zone rank x time: *F*=3.518, *p*<0.05, wait zone rank x time: *F*=103.712, *p*<0.0001) or for all groups splitting flavor rankings (right). **g** Standard deviation of offer zone and wait zone thresholds calculated across the four flavors within each 2.5 min bin (treatment x schedule: offer zone: *F*=56.443, *p*<0.0001, wait zone: *F*=9.031, *p*<0.01). **h** Standard deviation of the number of rewards earned calculated across the four flavor rankings within each 2.5 min bin (main effect of treatment: *F*=60.858, *p*<0.0001). Shading in (a,c) / error bars represent ±1 SEM.

By fitting decision thresholds restricted to sliding windows of time across the session, we found that VEH-treated mice displayed a fundamentally distinct decision-making profile compared to STZ-treated mice. Overall, in VEH-treated mice, offer zone thresholds significantly decreased across the session while wait zone thresholds remained relatively unchanged (Fig. 5e). Whereas in STZ-treated mice, the opposite and inverse was true: offer zone thresholds remained relatively unchanged while wait zone thresholds significantly increased (Fig. 5e). When segregating this analysis by restaurant, complex profiles emerged that were more dynamic with respect to flavor in VEH-compared to STZ-treated mice (Fig. 5f). In VEH-treated mice, offer zone thresholds for all flavors started at ∼30 s and decreased across the session, predominantly for less preferred flavors. Wait zone thresholds for all flavors started at ∼10 s and bifurcated across the session. In contrast, STZ-treated mice displayed initially flavor-disparate offer zone thresholds that were relatively stable across the session whereas wait zone thresholds, while also initially flav or-disparate, increased in all restaurants. These distinct profiles, summarized by taking the standard deviation of thresholds among restaurants across the session, were more different between VEH- and STZ-treated mice tested on the gradual than stepwise schedule (Fig. 5g). These data explain the decision-making policies behind meal consumption patterns across the session: VEH-treated mice focused first on securing food instead of exacting preferences by accepting all offers and waiting the optimal food-maximizing threshold in the wait zone, before subsequently laxing policies^21,27^; STZ-treated mice were skewed toward investing in more expensive offers to earn disproportionately-more higher-preferred flavors at the expense of yielding less food overall as a consequence (Fig. 5h, Supplementary Fig. 3i-p, recall Fig. 4e-f). These findings suggest STZ-altered sensitivity to economic choice competes with basic food security needs and depends on (i) the subjective value of the reward target at hand [flavor], (ii) the decision algorithm engaged [zone], and (iii) is influenced by the salience of reward scarcity in the environment derived from one’s prior history [gradual vs. stepwise].

### STZ-treated mice reveal diminished economic choice conflict

In addition to capturing economic conflict in the form of offer zone vs. wait zone decisi on policies, we measured a separate form of conflict within-trial during the decision process itself. We quantified behavioral path trajectories as mice traversed through the offer zone in route to making a skip or enter decision (Fig. 6a). Video-tracked body positions during the pass through the offer zone choice point can be transformed into absolute integrated angular velocity – a metric of hesitation or physical “hemming and hawing” known as vicarious trial and error (VTE) behavior^29-31^. VTE behavior has been previously shown to correlate with alternating neural representations of deliberation between competing choice options^32,33^. Overall, skip decisions elicited greater VTE behavior in the offer zone compared to enter decisions (Fig. 6b). The size of this discrepancy scaled with the ordinal ranking of flavor preferences (Fig. 6c, Supplementary Fig. 4). These data indicate mice demonstrated an aversion to skip in the offer zone that grows stronger with subjective flavor preferences. We discovered STZ-treated mice, particularly those previously tested on the gradual schedule, displayed diminished conflict that was more pronounced for less preferred rewards (Fig. 6c, Supplementary Fig. 4). We also found that this difference is related to but only partially explains the discrepancy between offer zone and wait zone thresholds (Supplementary Fig. 4). These data indicate that within-trial conflict during the choice process itself in the offer zone captures a unique aspect of reward valuation during planning behaviors altered by STZ treatment.

**Fig. 6.**
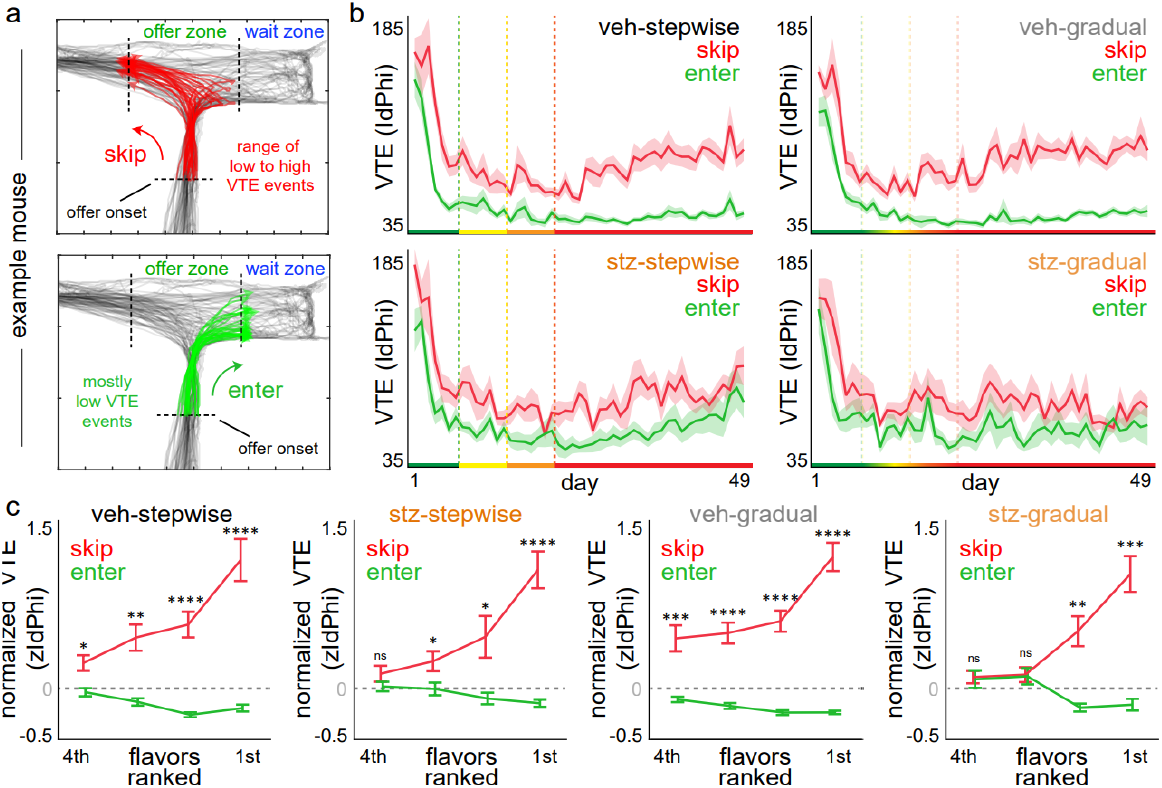
STZ-treated mice demonstrate diminished choice conflict when deliberating. **a** Example video-tracked body centroid positions in one restaurant from a single mouse on a single day. All tracked positions in black, skip decisions in red (top), enter decisions in green (bottom), tracked from offer onset upon entering the T-shaped choice point until crossing either boundary (skip: leftward hallway entry; enter: rightward wait zone entry). High vicarious trial and error (VTE) trials capture multiple reorientation events in the offer zone. **b** VTE behavior across the entire Restaurant Row paradigm (main effect of offer zone outcome skip > enter: *F*=121.070, *p*<0.0001). **c** VTE behavior, first normalized to all trials on a given day for a given mouse, then collapsed across the entire 1-30 s epoch and split by flavor rankings (offer zone outcome x rank: *F*=50.263, *p*<0.0001, treatment: *F*=5.876, *p*<0.05). Horizontal dashed gray line represents z-score of 0. Shading / error bars represent ±1 SEM.

### STZ-treated mice carry altered mental accounts of time

Next, we characterized the different ways in which mice value the passage of time on the Restaurant Row task. First, we examined ongoing choice processes in the wait zone by measuring the amount of time elapsed during the countdown before making a quit decision. When offer zone thresholds are higher than wait zone thresholds, this generally results in mice being more likely to quit accepted offers in the wait zone but does not capture how long it takes to quit. Overall, we found that all STZ-treated mice spent significantly more time waiting before deciding to quit compared to VEH-treated mice, regardless of prior schedule (Supplementary Fig. 5a,c). This was driven by trials in which STZ-treated mice accepted long-delay but not short-delay offers (Supplementary Fig. 5e-f). Furthermore, increased latencies to quit scaled with flavor in VEH-but not STZ-treated mice (Supplementary Fig. 5b,d). These data suggest that VEH-treated mice display an aversion to quit in the wait zone, like an aversion to skip in the offer zone, tied to subjective value but distinct from other forms of time spent on the task (Supplementary Fig. 5-7) and whose relationship is blunted in STZ-treated mice.

Why the majority of quit trials occur on long-delay offers can be due to the fact that mice have more of an opportunity to quit. However, as we have previously characterized, there is more value structure to these quit decisions that effectively correct offer zone mistakes in an economically advantageous manner (Supplementary Fig. 5k-o)^24,34^. What is more, quit events comprise a unique economic dilemma in which the longer an animal takes to decide to quit, the closer it may be to earning a reward. This highlights a conflict of deciding whether to continue waiting, encompassing a well-studied cognitive phenomenon known as the sunk cost bias. This describes the tendency to escalate commitment to an ongoing investment due to an accumulation of irrecoverable losses that, according to classic economic theory, should be ignored. We previously developed a dynamic analysis capable of extracting the influence of time spent in the wait zone on the likelihood of quitting and that is orthogonal to temporal distance to the goal^24,34^. First, each quit event was binned into [time-spent, time-left] pairs (Supplementary Fig. 5k). From this, we calculated the probability of earning a reward using a sliding-window survival analysis as animals continuously reevaluated staying in the wait zone (Fig. 7a-c, Supplementary Fig. 8a-f).

**Fig. 7.**
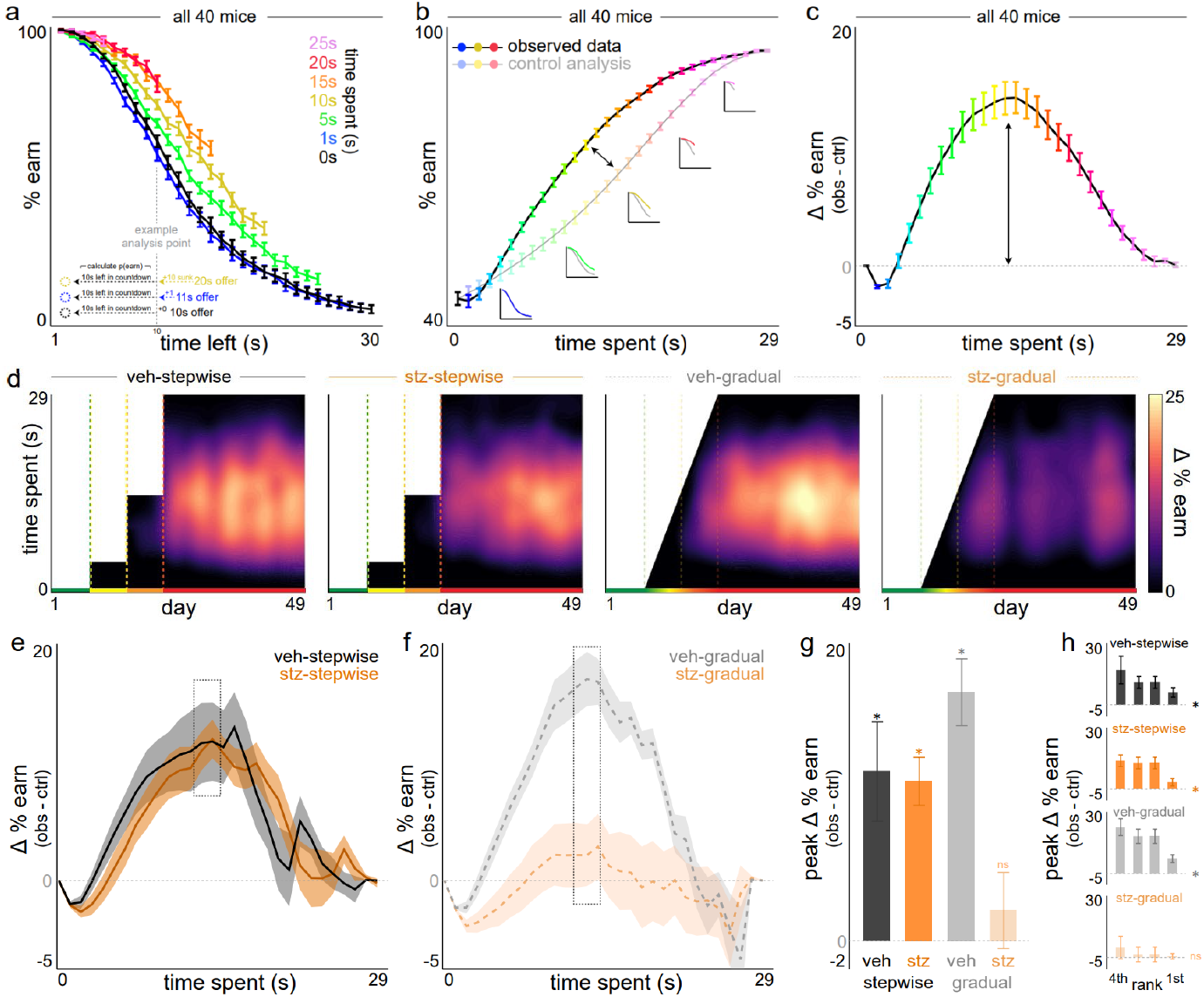
Only STZ-treated mice previously tested on the gradual schedule are uniquely insensitive to sunk costs. **a-c** Sunk cost analysis of staying behavior in the wait zone, demonstrated using all mice. (a) The likelihood of staying in the wait zone and earning a reward (e.g., not quit) is plotted as a function of time left in the countdown along the x-axis and time already spent waiting orthogonally in color. Note the black 0 s time spent curve represents animals having just entered the wait zone from the offer zone. Inset vertical dashed gray line illustrates an example analysis point comparing three sunk cost conditions originating from different starting offers but matched at 10 s left. Data from (a) dimensioned reduced in (b) collapsing across time left, instead highlighting the grand mean of each time spent sunk cost condition (color and x-axis). Insets depict data from curves in (a) are collapsed into the observed (sunk condition) and control (0 s condition) lines. Difference between curves in (b) are plotted in (c) in order to summarize the envelope of the overall effect of time already spent on escalating the commitment of staying in the wait zone. Horizontal dashed line represents 0. **d** Sensitivity to sunk costs plotted across the entire Restaurant Row paradigm where the delta curve in (c) is plotted as vertical slices in (d) with time already spent along the y-axis and heatmap representing magnitude. Note no sensitivity to sunk costs until after transitioning to the 1-30 s epoch in all mice, and never in STZ-gradual mice. **e-f** Delta curves collapsed across the entire 1-30s epoch for (e) stepwise or (f) gradual schedules (VEH: stepwise: *F*=76.544, *p*<0.0001; gradual: *F*=292.004, *p*<0.0001; STZ: stepwise: *F*=142.921, *p*<0.0001). Dashed gray box highlights peak differences. **g** Peaks from (e-f) for all groups (treatment x schedule: *F*=7.759, *p*<0.01). **h** Peaks split by flavor ranking. Shading / error bars represent ±1 SEM.

Consistent with our previous reports, we found that, in VEH-treated mice, the probability of earning a reward significantly increased as a function of time already waited (i.e., sunk costs, isolated from time left) but only after transitioning to the 1-30 s reward-scarce environment (Fig. 7d-e). In STZ-treated mice, we found that sensitivity to sunk costs depended on the history of their prior environment schedule. STZ-treated mice that previously experienced the stepwise schedule displayed robust sensitivity to sunk costs (Fig. 7d-e). However, STZ-treated mice tested previously on the gradual schedule displayed diminished sensitivity to su nk costs that was abolished across the entire experiment (Fig. 7d,f-h. Supplementary Fig. 8g).

Because sunk cost sensitivity emerges only after the transition to the 1-30 s reward-scarce environment indicates that this valuation process may depend on enhanced appetitive motivational levels driven by lower food availability. That is, provided individuals have intact systems capable of sensing this environmental change. VEH-treated mice remain attuned to this level of reward-scarcity regardless of prior schedule. And, while such a process may be impaired by STZ treatment, the stepwise schedule appears to preserve sunk cost valuations in STZ-treated mice conferred through the higher contrast in environmental change compared to the gradual schedule. Thus, we predicted that decreasing the relative scarcity of the environment by reducing food pressure ought to diminish sunk cost valuations as this may be a driver of blunted sensitivity in STZ-gradual mice. To test this without changing the distribution of offers in the environment, we restored *ad libitum* access to food in the home cage for all animals for an additional 5 days and continued restaurant testing. In addition to the expected decreases in laps run, pellets earned, and travel time (Fig. 8a-c, Supplementary Fig. 9a,d, Supplementary Fig. 10), we found differential changes in thresholds. While offer zone thresholds did not change in any group (Fig. 8d, nor offer zone VTE behavior, Fig. 8f), wait zone thresholds decreased in VEH-but increased in STZ-treated mice, regardless of prior schedule (Fig. 8e, Supplementary Fig. 9b). Lastly, we found that sensitivity to sunk costs remained intact in VEH-treated mice but was abolished in STZ-treated mice regardless of prior schedule experience (Fig. 8g, Supplementary Fig. 9e), reflecting a core phenotype of altered mental accounting unique to STZ animals.

**Fig. 8.**
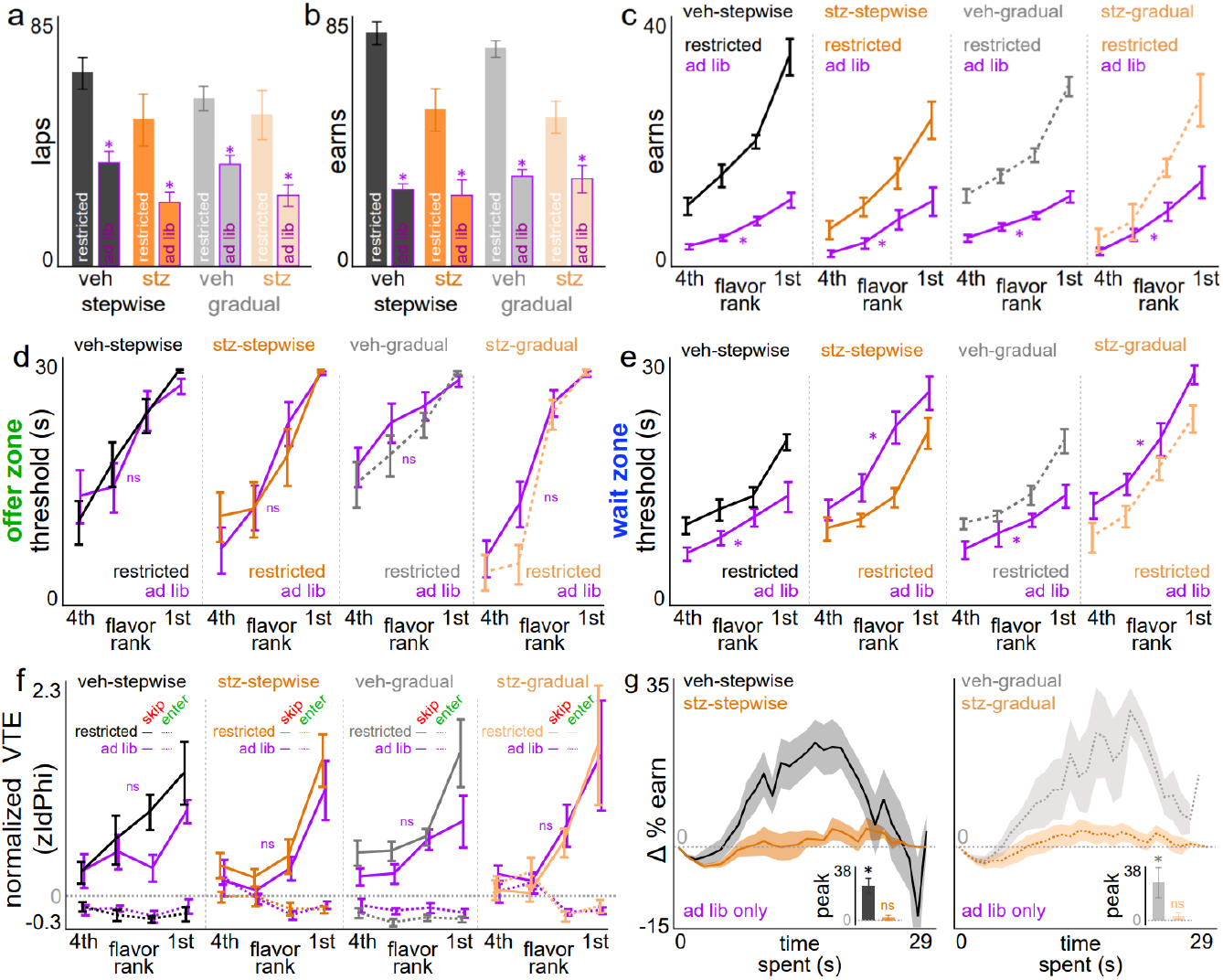
Restoring ad libitum access to food only affects re-evaluation decisions, differentially in VEH-vs. STZ-treated mice. **a-b** Number of laps run in the correct direction (a) and total rewards earned (b) comparing between when mice were food restricted vs. had unlimited access to regular chow in the home cage (purple outlines, main effect of ad lib vs. restricted: laps: *F*=41.383, *p*<0.0001; earns: *F*=45.597, *p*<0.0001). **c** Rewards earned split by flavor rankings. **d-e** Offer zone thresholds (d) and wait zone thresholds (e) split by flavor ranking. Note no change in offer zone thresholds (*F*=1.725, *p*=0.190) but a bidirectional change in wait zone thresholds (*F*=21.027, *p*<0.0001, VEH: decreased; STZ: increased). **f** Normalized vicarious trial and error (VTE) behavior split by flavor ranking and by skip vs. enter offer zone outcomes (no effect of treatment on enter: *F*=0.346, *p*=0.557, or skip: *F*=0.791, *p*=0.375). **g** Sensitivity to sunk cost delta curves plotted as a function of time already spent in the wait zone. Note abolished sunk costs in both STZ-treated groups regardless of schedule (main effect of treatment: *F*=15.045, *p*<0.001), in spite of no change in offer zone thresholds and similar shifts in wait zone thresholds. Insets depict peak scores. Horizontal dashed gray line represents 0. Shading / error bars represent ±1 SEM.

## Discussion

We characterized complex neuroeconomic decision-making behavior on the Restaurant Row paradigm in chronically hyperglycemic mice using the STZ diabetic mouse model. We tested mice longitudinally across months in a closed-economy system. We found that mice learned to respond to the changing economic landscape of rewards distributed throughout the environment as a function of cost in the form of delays required to earn food and subjective value in the form of flavors. We discovered diabetic mice employed altered economic decision-making policies that interacted with revealed preferences, manifested in only certain types of choices, and influenced the value of the passage of only certain categories of time. We also discovered that the effects of diabetes on neuroeconomic valuation processes depended on one’s prior experience with how rapidly the environment changed from reward-rich to reward-scarce, and were more pronounced when this transition was gradual. These findings suggest that diabetes is capable of altering dissociable forms of mental accounting as a result of a complex interaction between energy demand, multiple decision-making algorithms accessible to the agent, and the salience of scarcity of reward availability in the environment.

Diabetes mellitus is characterized by dysregulated glucose metabolism mediated through impairments in insulin signaling either as a consequence of deficient insulin production or insulin receptor resistance^1,35^. How diabetes may affect motivated behavior is multifactorial. Principally, increased circulating levels of glucose in the bloodstream and lower levels of glucose uptake in tissue throughout the body is capable of altering peripheral drivers of reward-seeking behavior as a result of changes in global energy levels, body muscle and fat composition, and bottom-up signals of hunger and satiety^2^. As a key example, inability to utilize glucose as a peripheral fuel-source leads to fatty-acid oxidation and weight loss in STZ-treated rodents, with a subsequent reduction in circulating leptin levels that are thought to contribute to the hyperphagia associated with diabetes via a central mechanism^36,37^. Similar dysregulation of other hormones (e.g. cortisol, ghrelin) also likely contribute to altered reward behavior in diabetes^38-40^. Centrally, elevated circulating levels of glucose in the brain lead to reduced cerebral perfusion as means to limit more glucose entry^2^. This results in decreased blood flow and oxygen delivery that could interfere with ongoing cognitive processes, which may be contributing to differences in motivation or altered decision-making in diabetes^5,41^. Reduced processing speeds on simple cognitive tasks have been reported in people with diabetes but with little done thoroughly characterizing aspects of reward and motivation in either human or animal studies^3,6^. Indeed, diabetes in animals can impair novel spatial and object recognition as well as performance in Barnes, Morris water-maze, and active-avoidance foot-shock tasks, aspects of which may be correctable with glycemic control^42-44^. However, glucose is not the only fuel available to the brain; the brain can also utilize ketones as a fuel source, although this switch in metabolism requires mitochondrial adaptations, some of which have been linked to changes in motivated behavior^2,45,46^. More recent work has implicated glucose-independent roles of insulin function directly in the central nervous system^10-14^. Insulin is actively transported across the blood-brain barrier and has been shown to alter the function of the mesolimbic dopamine system^14^. Insulin receptors are present on both ventral tegmental area dopaminergic neurons, as well as on neurons in the nucleus accumbens, where activation of the receptor by insulin is thought to suppress hedonic feeding^35,47-49^. The observation that STZ-mice are biased for most-preferred flavors is consistent with a role for insulin in hedonic feeding. Thus, the absence of central insulin in Type 1 diabetes may have profound consequences on dopaminergic function, either acutely by altering transient activity or also across longer timescales affecting plasticity and reward circuit remodeling. Such changes could influence not only reward-related behavior during meals but also impact decision-making information processing more broadly, giving rise to complex psychiatric disease vulnerabilities^35,47^. Given the more complex and multifactorial changes in dissociable decision-making behaviors observed on the Restaurant Row task in diabetic mice, we can begin to point to multiple, parallel circuit-computation-specific valuation algorithms that may be uniquely disrupted in diabetes.

Separate behaviors in the offer zone and wait zone reflect fundamentally distinct types of choices^22,50^. In the offer zone, mice are faced with choosing between accepting versus rejecting an offer whose cost is cued, but an investment has not yet taken place. Whereas in the wait zone, mice are continuously re-evaluating commitment to an ongoing investment as they temporally approach earning a reward. We and others previously demonstrated that offer zone decisions and wait zone decisions access computationally distinct functions of physically separable circuits in the brain^50,51^. The cognitive mechanisms underlying offer zone behaviors have been previously linked to deliberative decision-making processes shown to engage circuits that support prospective thinking^52,53^. Hippocampal recordings during such choices are capable of decoding alternating representations of competing actions that sweep ahead of the animal through the choice point leading to potential future goal locations^29-33^. Failure to engage in such processes can cause individuals to rely on other Pavlovian or procedural memory systems^18,27,54-57^. Chemogenetically inactivating the medial prefrontal cortex can disrupt synchrony with the hippocampus, impair hippocampal sequences in the offer zone, and reduce deliberative behaviors such as VTE^52,56^. In humans tested on translated variants of the Restaurant Row task, functional magnetic resonance imaging of the default mode network, including hippocampal and prefrontal regions, has been shown to decode current and future goal locations prior to making decisions in the offer zone^58,59^. Conversely, the cognitive processes underlying wait zone behaviors are thought to capture change-of-mind decisions that have far less understood neural mechanisms^60,61^. Continuously re-evaluating an ongoing investment on Restaurant Row encompasses a complex integration of requiring mice to patiently wait in a confined location while counting the passage of time^34^. With this, animals are losing irrecoverable time, have more of a window to reconsider opportunity costs in tandem with growing hunger states, all the while decreasing temporal distance to the goal. Recent recordings of dopamine release in the striatum using fiber photometry during Restaurant Row have revealed differential signaling patterns of costs coded in the offer zone versus wait zone^62^. Further, dopamine encoded perplexing negative reward-prediction-error-like signals time-locked to volitional change-of-mind quit decisions in the wait zone that could be disrupted optogenetically^62,63^. Altering synaptic strength of glutamatergic inputs into the striatum can also selectively disrupt wait zone behavior without influencing offer zone choices^64^. Together, the changes observed in offer zone and wait zone thresholds, VTE behavior, and wait zone quitting behavior of STZ-treated mice characterize a suite of alterations in information processing that moves beyond simple dysfunction of central glucose metabolism and suggests a more complex and dynamic dysregulation of multiple decision-making systems in diabetes.

In addition to understanding complex decision systems in the brain, interactions between an agent and one’s changing environment is critically important^65^. We found that, by and large, most of the differences in decision-making behavior between STZ- and VEH-treated mice emerged only in those tested across an environment that grew increasingly reward-scarce on a gradual but not stepwise schedule. We previously demonstrated that the abruptness of the stepwise schedule serves as a naturalistic economic stressor capable of interacting with one’s prior history of stress and can alter subsequent decision-making strategies^21,66^. Thus, the gradual schedule may serve as a control for such a striking economic stressor and could dampen the perception of reward-scarcity in the environment compared to the stepwise schedule, despite ultimately arriving in the same environment. This concept captures the well-known economic framework of creeping normality, also known as the allegory of the “boiling frog,” in which behavioral responses are blunted when external challenges escalate gradually^67^. The stepwise schedule may be triggering a switch into an altered brain-body state that could pressure the mobilization of energy stores or activate alternative decision-making strategies in VEH- and STZ-treated mice alike. However, the gradual schedule may elicit a sub-threshold economic stress response phenotype that when combined with diabetic physiology reveals striking differences in foraging behavior that are otherwise masked when environmental circumstances change abruptly. This could be in part due to impaired glucose sensing in diabetes that is dependent on the salience of or contrast in reward availability in the environment – a dysfunction exaggerated on the gradual schedule. This could also stem from decision-making algorithms that should switch in response to a changing environment, regardless of rate, but fail to do so in STZ-gradual mice only. Interestingly, we found that when placed back on ad lib access to food, despite observing changes in foraging that would be expected when sated in all animals (e.g., decreased laps run and pellets earned), specific decision-making phenotypes that may be a core feature of diabetes emerged. Most notably, this manifested as abolished sensitivity to sunk costs in STZ-treated mice irrespective of prior testing schedule.

Mice, like humans, are capable of valuing the passage of time in such a way that can inflate the value of continuing to invest in an ongoing endeavor^24,34^. This reflects the well-studied decision-making phenomenon termed the sunk cost fallacy, that according to classic economic theory, should be ignored when making re-evaluative choices^68^. We previously discovered that this valuation algorithm has been conserved across evolution postulated to arise from, among several reasons, state-dependent valuation learning as a key driver of this phenomenon that may be useful from an energetic standpoint^24,34,69,70^. This theory hypothesizes that energy expended in pursuit of a food reward could shift an individual into a poorer energy state^69,70^. As a result, the yet-to-be-earned food reward may have an enhanced perceived value that can escalate the commitment of continued investments^69,70^. Practically, this may help the agent in times of survival. We and others have observed sensitivity to sunk costs only in a reward-scarce environment, suggesting that this within-trial state dependent valuation learning explanation of sunk costs depends on task demand^21,24,34,71,72^. Thus, it appears that diabetic mice tested on the gradual schedule who fail to demonstrate sensitivity to sunk costs may do so in part because of a decreased pressure to seek food secondary to testing history or because of a blunted ability to register internal states with external circumstances. Nonetheless, this phenotype was further abolished when STZ-treated mice were placed back on ad lib food access unlike VEH-treated mice who maintained intact sensitivity to sunk costs, suggesting that this form of mental accounting may be uniquely perturbed in diabetes at its core.

Limitations of the present study include no direct measures of metabolic physiology, body composition, or calorimetry as it relates to energy expenditure of animals tested across this longitudinal experiment. Further, food restriction-based studies in diabetic animals are incredibly difficult to manage while ensuring animal safety, particularly at such long timescales. Future work should explore how behaviors shift under different economic constraints: for instance, (i) if mice have, instead of a fixed time-budget, a fixed number of choices with an open-ended amount of time; or (ii) instead of titrating mice to a fixed body weight, evaluating the weight individual mice settle at as a dependent outcome measure as a function of distribution of costs in the environment. After validating and eliciting robust diabetic phenotypes in female mice, which has limitations presently in the STZ model^26^, sex differences in the effects of hyperglycemia on decision-making behaviors should be explored next. Investigating the role of insulin signaling *in vivo* will be critically important in future studies, including comparing and contrasting the effects of Type 1 versus Type 2 diabetes on decision-making. This should take the form of experiments studying insulin replacement challenges but also measuring how peripheral and central targets of insulin might change throughout these two testing schedules or across different decision-making processes, including recording the activity of neural signals such as dopamine in diabetic mice on the Restaurant Row task.

We demonstrated that in a diabetic mouse model, complex neuroeconomic decision-making behaviors that are multifaceted and reflect fundamentally distinct valuation algorithms can be uniquely perturbed not only as a function of hyperglycemia but also as a consequence of environmental dynamics. We found diabetes-induced changes in how mice deliberate for and re-evaluate rewards of varying costs and subjective value in a manner that depends on the salience of reward-scarcity in the environment. Our task, which has been translated for use across species in humans^24,34,58,59,73,74^, allows for a thorough neuroeconomic investigation linking the biological mechanisms underlying metabolic disorders to unique decision-making vulnerabilities that could predispose individuals to develop co-morbid psychiatric illnesses.

## Acknowledgments

We thank members of the labs of Eric Nestler and Scott Russo for helpful discussion. Open-source illustrations obtained from SciDraw (www.scidraw.io), credit Federico Claudi.

## Funding

National Institute of Mental Health grant L40MH127601 (BMS)

National Institute of Mental Health supplement grant R01MH051399-31S1 (BMS)

Leon Levy Scholarship in Neuroscience, New York Academy of Sciences (BMS)

Burroughs Wellcome Fund Career Award for Medical Scientists (BMS)

Animal Models for the Social Dimensions of Health and Aging Research Network via NIH/NIA R24 AG065172 (BMS)

Brain & Behavior Research Foundation Young Investigator Award 31140 (RDC)

Brain & Behavior Research Foundation NARSAD Young Investigator Award 28240 (JLA)

Einstein-Mount Sinai Diabetes Research Center Pilot & Feasibility Award P30DK020541-FS553 (JLA)

Mount Sinai SURP4US (CAN)

## Author contributions

Conceptualization: JLA, BMS

Methodology: JLA, BMS

Investigation: CAN, RDC, ZMO, SOB, JEH, AM, MJF, YZC, SA, SL, JLA, BMS

Data curation: CAN, RDC, JLA, BMS

Formal analysis: CAN, RDC, JLA, BMS

Visualization: CAN, RDC, BMS

Funding acquisition: JLA, BMS

Supervision: RDC, JLA, BMS

Writing – original draft: CAN, JLA, BMS\

Writing – review & editing: all authors

## Data and materials availability

All data, code, and materials used in the analysis are available in the manuscript, materials and methods section, supplementary information, or upon request.

## Methods

### Subjects

Adult male C57BL/6J mice (Jackson Labs, stock # 000664) were used in this study. Mice were maintained on a 12-hr regular light/dark cycle with ad libitum access to water throughout the entire experiment. During the treatment protocol to induce chronic hyperglycemia, mice were group-housed with ad libitum access to regular chow (LabDiet 5053; Protein 21%, Fat 6%, Carbohydrate 53.5%, Fiber 4.4%, Ash 6%, 4.11 kcal/gm). Beginning 3d prior to and continuing during testing on the Restaurant Row paradigm, mice were single-housed to avoid shared flavor preferences and food restricted to 80-85% of pre-task body weight. Behavioral testing was conducted during the light phase in dim lighting. Experiments were approved by the Mount Sinai Institutional Animal Care and Use Committee (IACUC; protocol number LA12-00051) and adhered to the National Institutes of Health (NIH) guidelines.

### Chronic Hyperglycemia

At 8 weeks of age, 40 C57BL/6J mice received a daily intraperitoneal injection of either Hank’s balanced saline solution (HBSS) as the vehicle (VEH) control or streptozotocin (STZ, 50 mg/kg), an antibiotic that ablates insulin producing beta cells in the pancreas, for 5 consecutive days to induce hyperglycemia (Fig. 1a)^26^. Mice were maintained on ad libitum food for the next 6 weeks to allow for assessment of chronic hyperglycemia. Body weights and fasting blood sugar (morning fast, ∼5 hrs) were assessed weekly to monitor progression of diabetes using a handheld glucometer (Bayer Contour). At 14-weeks of age, Hemoglobin A1C (HbA1c) tests of tail vein blood were performed to confirm a hyperglycemia phenotype using A1CNow+ point-of-care kits (PTS Diagnostics). Mice were then single-housed, and food restricted to 80-85% of their free-feeding body weight over the 3 days preceding the beginning of the task.

### Neuroeconomic Decision-Making Paradigm

20 VEH- and 20 STZ-treated mice were then characterized longitudinally on the Restaurant Row task^27^. On this task, mice had a limited amount time each day to forage for their primary source of food by navigating a maze with four uniquely flavored and contextualized feeding sites, or “restaurants” (Fig. 2b). Food rewards consisted of 20 mg full-nutrition pellets (BioServ dustless precision pellets) that varied in flavor (chocolate, banana, grape, or plain) but not caloric content (3.6 kcal/gm calories). Macronutrient content did not vary grossly across flavors (18.7% protein, 5.6% fat, 59.1% carbohydrate, 4.7% fiber, 6.5% ash for banana, grape or plain; 18.4% protein, 5.5% fat, 59.1% carbohydrate, 4.6% fiber, 6.5% ash for chocolate). Each restaurant was decorated with either horizontal stripes (chocolate), dots (banana), triangles (grape), or vertical stripes (plain), whose locations remained spatially fixed throughout the entire paradigm. Animals were placed into an arena (approximately 36” x 36”) consisting of 4 fixed restaurants positioned in the corners of a square maze and connected by hallways for 45 min (their limited daily time budget) and video-tracked under dim lighting using a standard USB camera and computer running the task programmed in ANY-Maze (Stoelting). Animal position tracking in real-time controlled engagement with the task whereby centroid body crossings into distinct zones in the maze triggered task events, including speaker playback of various tones or activating custom-built 3D-printed automated pellet dispensers (open-source pellet dispensers used in this experiment: www.hackaday.io/project/171116-fed0). Task rules required animals to run in a counterclockwise direction so that they properly approached each restaurant’s T-shaped intersection in order engage task contingencies and trigger events, which all animals quickly acquired within the first week of testing. Each restaurant had a separate offer zone and wait zone. Upon entry into the offer zone from the correct heading direction, a tone sounded whose pitch indicated the offer length of how long of a delay mice would have to wait in a cued countdown should they choose to enter the wait zone in order to earn a food reward. The range of tone pitches used in this paradigm varied from 4,000 Hz (lowest pitch, signaling a 1 s offer) in increments of 387 Hz to 15,223 Hz (highest pitch, signaling a 30 s offer), determined by the schedule animals were assigned to, described in detail below. In the offer zone, tones played for 500 ms and repeated every second at the same pitch until either an enter decision was made by turning right into the wait zone or a skip decision was made by turning left into the hallway advancing toward the next restaurant. If mice choose to enter the wait zone, tones (500 ms) descended by 387 Hz every second signaling the countdown to reward delivery. If mice decided to quit and exit the wait zone prematurely before the countdown completed, tones silenced, the offer was rescinded, and mice must advance to the next trial in the next restaurant. Thus, this task is economic in nature, requiring animals to budget their limited time effectively in a self-paced manner in order to earn a sufficient amount of food. Animals were maintained at 80-85% body weight per IACUC regulations and both VEH and STZ groups were supplemented with additional food post-task if needed to maintain safety.

A key manipulation and focus of the present study was the way in which the distribution of offers presented to the animals changed across weeks. For the first week of behavioral testing, all offers were 1 s only (i.e., the lowest 4,000 Hz pitch tone presented, green epoch). This allowed animals to easily learn the basic structure of the task and reveals individual differences in flavor preferences. Beginning on day 8 until day 22, the range of offers presented to the animals increased, ultimately to the final 1-30 s offer range, but did so via one of two schedules: (i) stepwise or (ii) gradual. For the stepwise schedule, during the next 7 days of testing (block 2, days 8-14, yellow epoch), offers ranged from 1 to 5 s; block 3 (days 15-21, orange epoch) consisted of a 1 to 15 s range. The fourth and final block (days 21-49, red epoch) consisted of offers ranging from 1 to 30 s. For the gradual schedule, during the next 15 days of testing (days 8-22), the maximal offers increased each day by 2s (e.g., day 8: 1-2 s range; day 9: 1-4 s range; day 10: 1-6s range, and so on) until reaching the same 1-30 s offer range by day 22. These two schedules allowed us to examine how mice initially shaped their behavior as they learned the basic structure of the task as well as how they adjusted their foraging strategies across long timescales (days, weeks, and months) as they transitioned from reward-rich to reward-scarce environments in either a stepwise or gradual manner. This design optimizes the experimental plan, minimizing as many discrepancies between the schedules as possible. Regardless of the range of offers on any given day, offers were always sampled from a uniform distribution separately in each restaurant. Thus, after day 7, VEH- and STZ-treated mice were split to continue either on the stepwise or gradual schedule, randomly assigned and counterbalanced across numerous behavioral metrics. Animals developed strategies longitudinally across the changing economic landscape. Testing continued on the 1-30 s offer range until the main experiment terminated on day 49. At the very end of the experiment, after the main behavioral period during days 1-49 concluded, subjects were placed back on ad lib access to regular chow in the home cage for an additional 5 days while continuing testing on Restaurant Row in their final week. For graphical comparison, restricted baseline data either was calculated in the 2 days leading up to the ab lib sessions or replotted from the average of the entire 1-30 s epoch, indicated within each figure. Pre- and post-task tail vein blood glucose levels were sampled on days 7 and 22 in both the VEH- and STZ-treated animals. In the final week of testing, only the STZ-treated mice were sampled again for pre-task tail vein blood glucose levels to ensure safety and maintenance of hyperglycemia at long time points. Terminal HbA1c levels were measured in all mice at the conclusion of the study.

### Data & statistical analyses

Data were processed in Matlab with statistical analyses in JMP Pro 16. All data are expressed as mean ± 1 standard error. Statistical significance was assessed using student’s t tests and one-way, two-way, and repeated measures ANOVAs, and sign-tests. Correlations were reported using Pearson correlation r coefficients. Several behavioral analyses were developed and either previously published or newly described in this manuscript, detailed below. Many analyses of interest were calculated by collapsing across the entire 1-30 s epoch (days 22-49) unless otherwise specified.

Meal consumption patterns were calculated by binning the full 45 min session into 18 segmented 2.5 min bins. Within each bin, we calculated the total number of rewards earned, rewards earned in each restaurant, as well as cumulative sums of both metrics either in raw pellet counts or as a percentage of either total global earns or total earns within-flavor. This allowed us to capture rate of within-flavor meal consumption patterns. Flavors were ranked from most to least preferred each day by summing the end of session earn totals in each restaurant. To summarize mid-session meal patterns, we extracted % cumulative earnings within-flavor metrics at the half-way point, and calculated a difference score between the most preferred and least preferred flavors in this metric to generate a summary statistic measure reflecting a sign change in the temporal rank order of flavor-specific meal consumption.

Percent change in earns across the entire study was measured within-subject relative to the average of stable total earns obtained from days 5-7. This allowed us to capture, within-subject, how the subsequent changing economic landscape of the task caused a decrease in total earnings relative to this day 5-7 average (termed 100%). From this, we calculated difference scores in this metric at each transition point in the stepwise schedule (i.e., day 7-8, day 14-15, and day 21-22) showing the drastic stair step effects on earnings this schedule has on both VEH- and STZ-treated mice compared to the gradual schedule.

In order to approximate economic thresholds, or indifference points, of willingness to accept and earn rewards, we fit a heaviside-step regression to choice outcomes as a function of cost in each zone and measured curve inflection points each day across the entire experiment. Wait zone thresholds, which determine offers animals are actually willing to wait for and earn and are relatively stable, are used in calculations to determine the value of an offer on any given trial for that day (offer value = wait zone threshold minus offer). This allows offers to be normalized into value terms across animals, across days, or within-animal across flavors. Thus, thresholds were recalculated using a leave-one-out method on a trial-by-trial basis to avoid conflating value normalization with the offer presented and behavioral outcome on any given trial. The value remaining at the moment of quitting could similarly be calculated (value left = wait zone threshold minus time remaining in the countdown at the moment of quitting) to categorize quit events into separate economic categories. Difference scores were also calculated between offer zone minus wait zone thresholds in order to summarize to what degree thresholds were in register (0) or out of register with one another (e.g., thus yielding quitting behaviors in the wait zone).

In order to capture deliberation behaviors, we quantified behavioral path trajectories as mice traversed through the offer zone in route to making a skip or enter decision. Video-tracked body positions during the pass through the offer zone choice point can be transformed into absolute integrated angular velocity – a metric of hesitation or physical “hemming and hawing” known as vicarious trial and error (VTE) behavior. Trajectories in this analysis started at offer onset entering the stem of the T-shaped 180-degree choice point until either hallway entry (left turn, skip) or wait zone entry (right turn, enter). This behavior is best measured by calculating changes in velocity vectors of discrete body *x* and *y* positions over time as *dx* and *dy*. From this, we can calculate the momentary change in angle, *Phi*, as *dPhi*. When this metric is integrated over the duration of the pass through the offer zone, VTE is measured as the absolute integrated angular velocity, or *IdPhi*. For a given animal on a given day, we can normalize all *IdPhi* metrics obtained on every trial and generate a *zIdPhi* value that can then be split post-hoc for comparisons under any number of trial conditions (e.g., skip vs. enter, by flavor, etc). This allowed us to capture within-subject but between-decision-condition differences in internal choice patterns. Thus, we could calculate differences scores between *zIdPhi* when skipping minus entering, for instance as a within-subject metric of choice conflict.

In order to capture sensitivity to sunk costs, we previously developed a dynamic analysis capable of extracting the influence of time spent in the wait zone on the likelihood of quitting that is orthogonal to temporal distance to the goal. First, each quit event was binned into [time spent, time left] pairs. From this, we calculated the probability of earning a reward using a sliding window survival analysis as animals continuously reevaluated staying in the wait zone. In order to reduce the dimensions of this analysis, we collapsed each time spent condition across the time left axis. To control for artificial inflation of the probability of earning due to increasingly non-existent data being left out of the grand means at higher time spent conditions, we resampled data from the 0 s time spent condition and collapsed along the time left dimension iteratively excluding points to match each higher time spent condition. This resultant dimension-reduced analysis yields an observed and control curve that can be subtracted to yield the final delta curve capturing the envelope of sensitivity to sunk costs in altering staying behavior in the wait zone that is relative to each animal’s 0 s sunk psychometric staying function. Note the rising phase of this curve captures sensitivity while the falling phase of this curve diminishes due to ceiling effects of data samples used at high time spent conditions. Peaks were extracted from this delta curve for summary statistical comparisons between groups and tested against 0, or no sensitivity to sunk costs. Additional analyses of the effects of different forms of time spent on the probability of staying in the wait zone during the countdown, in contrast to sunk costs accrued in the wait zone, were also examined, including time spent deliberating in the offer zone before accepting an offer or time elapsed since last reward earned before accepting an offer. In both of such cases, the read-out metric of the probability of earning once in the wait zone was calculated.

Post-consumption behaviors were measured as time spent consuming and lingering at the feeding site after earning a reward from reward delivery onset until exiting the wait zone. Transit times between restaurants were measured from trial termination (i.e., skip onset, quit onset, or p ost-earn wait zone exit onset) until mice arrived at the next restaurant’s offer zone choice point, triggering the next trial onset. Inter-earn-intervals were calculated as the amount of time elapsed between subsequent earns either of any flavor or specifically of a same, given flavor.

## SUPPLEMENTARY INFORMATION

**Supplementary Fig. 1.**
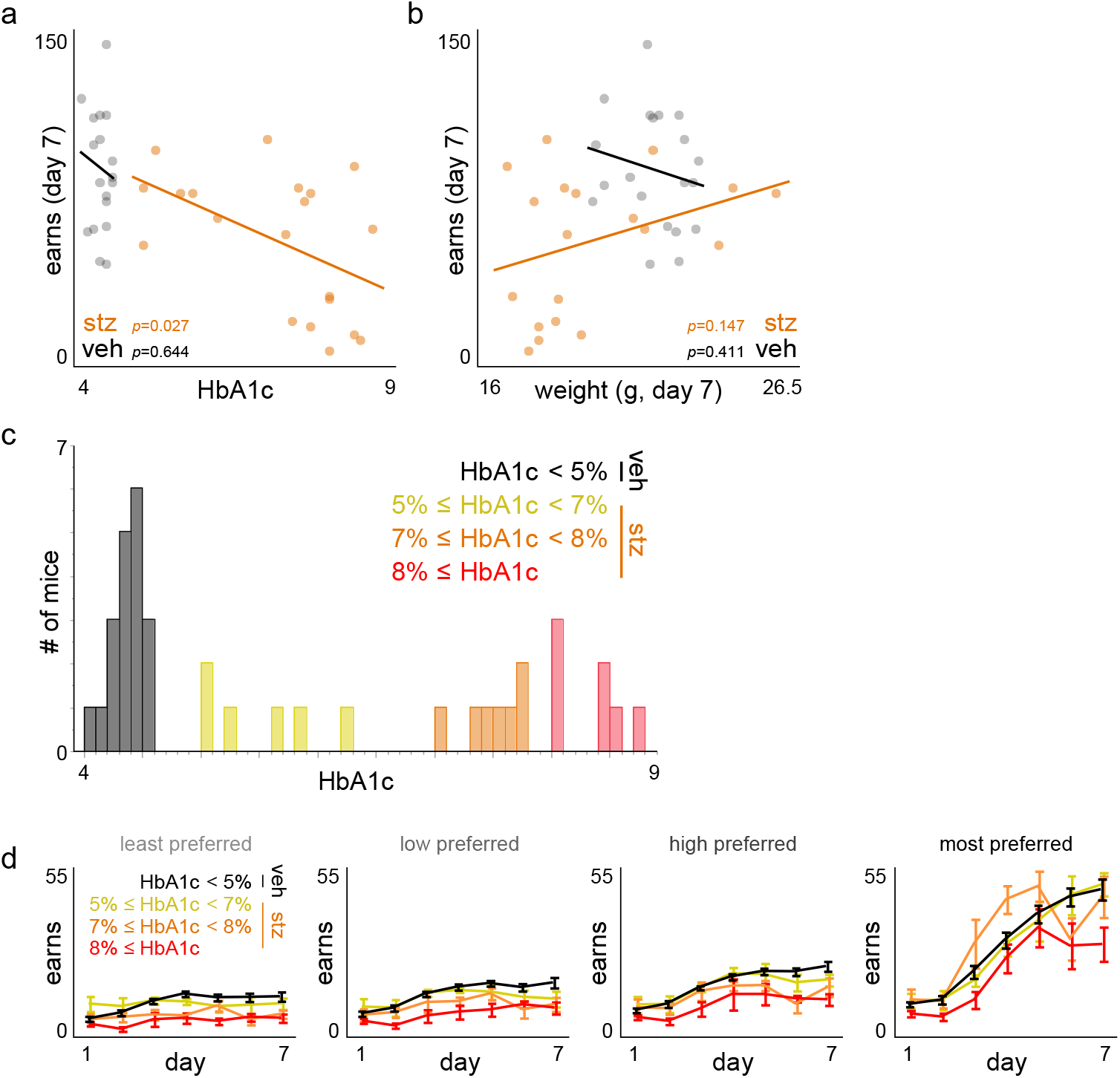
Analysis of HbA1c levels and behavior. **a-b** Scatter plot of (a) HbA1c levels or (b) pre-task body weight measured on day 7 immediately before behavioral testing against total rewards earned on day 7. Trendline represents linear fit. **c** HbA1c levels split into evenly distributed sub-groups. **d** Visualization of rewards earned across days 1-7 split by flavor rankings in each HbA1c sub-group. Dots represent individual mice. Error bars represent ±1 SEM.

**Supplementary Fig. 2.**
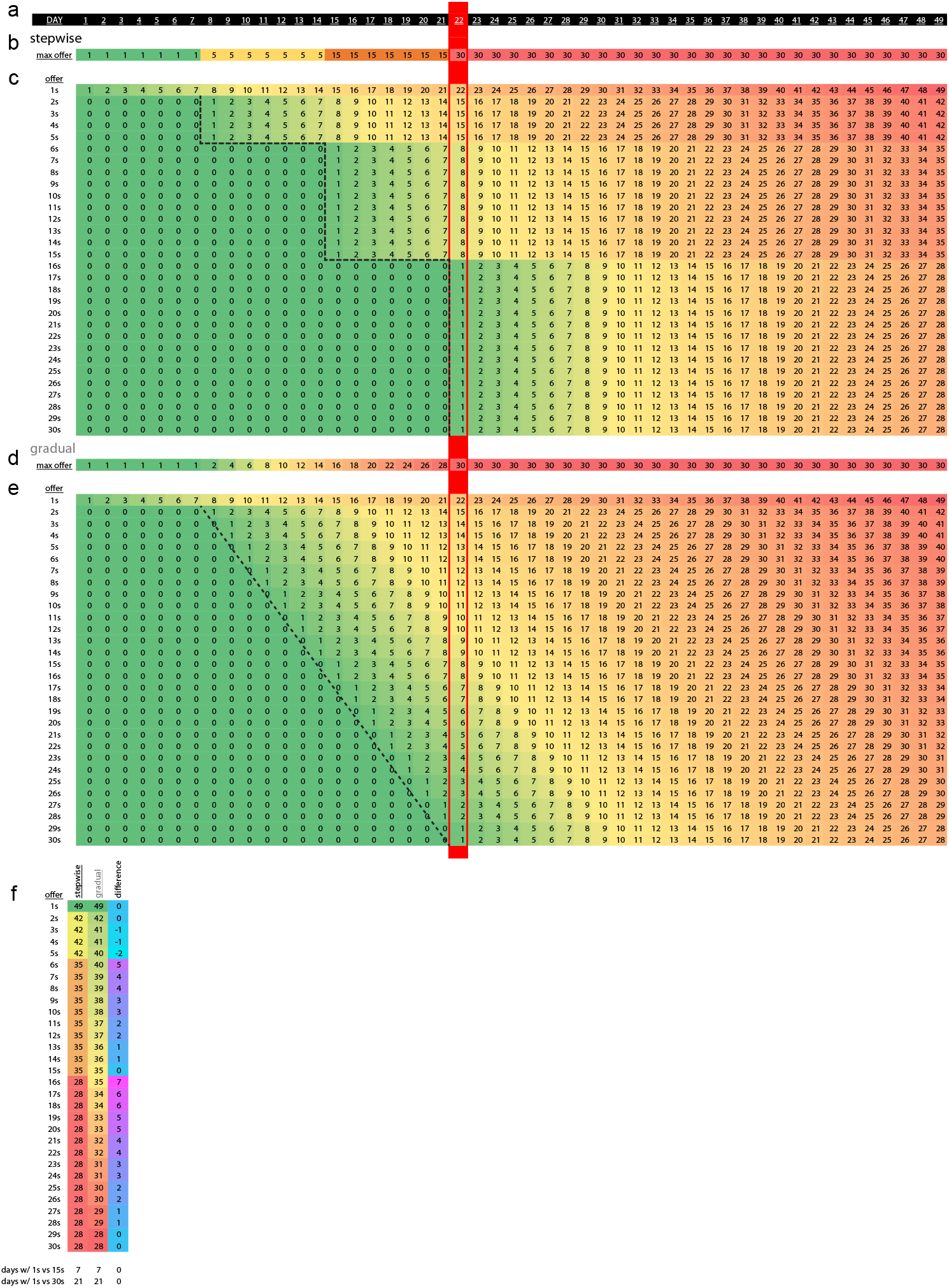
Visualization of experimentally manipulated Restaurant Row testing schedules. **a** Timeline of the entire Restaurant Row paradigm, days 1 to 49 enumerated. Red marking on day 22 indicates the first day of the transition into the 1-30 s reward-scarce epoch, matched for both schedules. **b** Stepwise schedule timeline. Max offer on each day enumerated and color coded. **c** Stepwise schedule visualization of cumulative number of days animals would have had experience with each offer length, enumerated and color-coded scale ranging from 0 days of experience (green) to 49 days of experience (red). Staircase dashed line represents transition points. **d** Gradual schedule timeline. Max offer on each day enumerated and color coded. **e** Gradual schedule visualization of cumulative number of days animals would have had experience with each offer length, enumerated and color-coded scale ranging from 0 days of experience (green) to 49 days of experience (red). Diagonal dashed line represents gradual transition points. **f** Comparison of metrics between stepwise and gradual schedules. Columns 1 and 2 enumerate total number of days of experience with each offer length. Column 3 represents the difference in total number of days of experience with each offer length between stepwise minus gradual schedules (cool color-code). Bottom two rows of table indicate the difference between number of days of experience with 1 s offers vs 15 s offers or 1 s offers vs 30 s offers between the stepwise and gradual schedules, which are matched. This experimental design optimally controls for as many differences between schedules while varying rate of environmental change.

**Supplementary Fig. 3.**
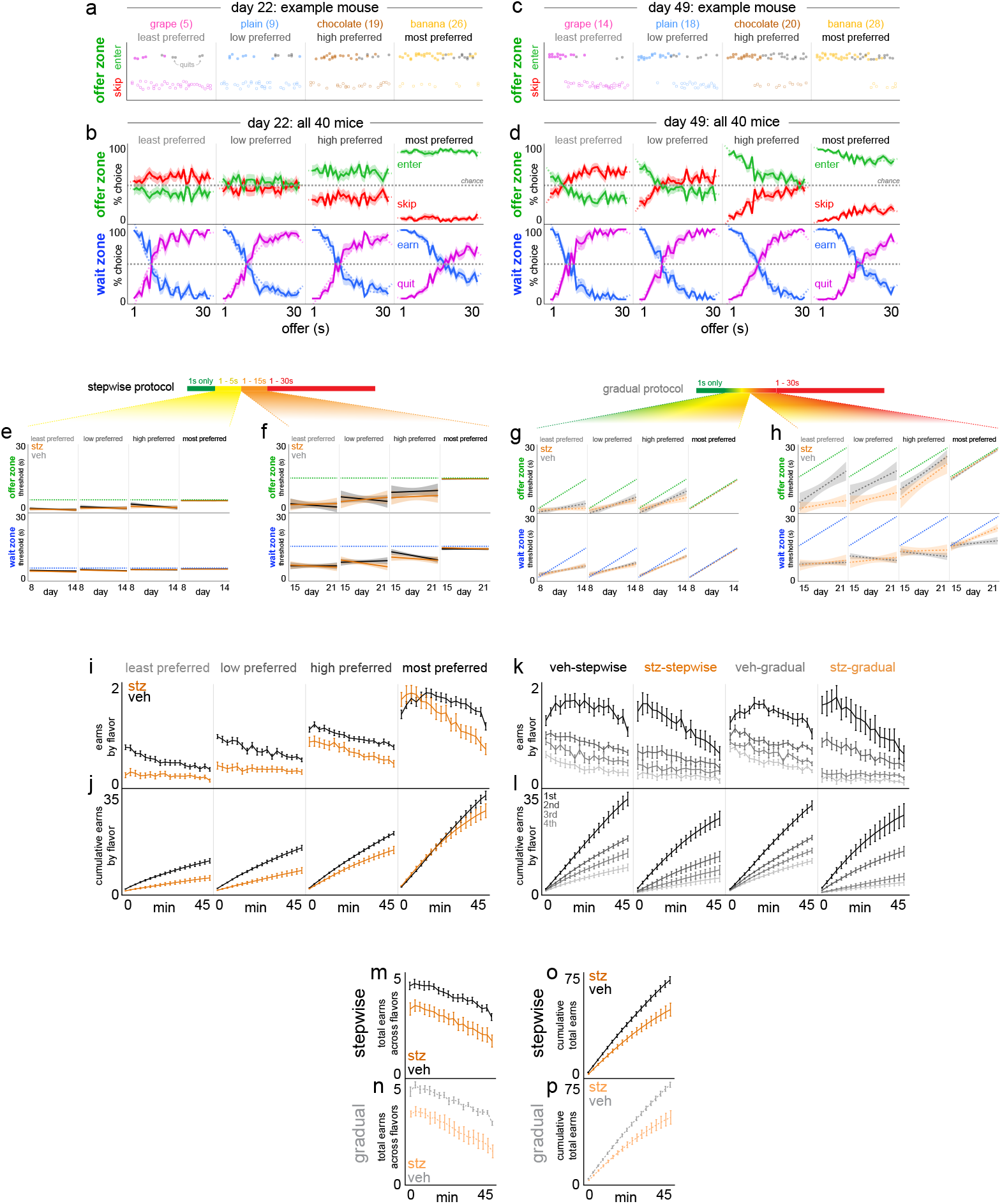
Expansion of choice behavior metrics on Restaurant Row. **a-d** Example behavior from a single mouse on (a) day 22 and (c) day 49 as well as aggregate example summary choice data from all 40 mice on (b) day 22 and (d) day 49. Data depict offer zone choice outcomes on the y-axis (enter vs. skip) as a function of cued offer cost (tone pitch) on the x-axis either as individual choices (dots in a,c) or as summary % choice (curves in b,d). Horizontal dashed gray line in (b,d) indicates chance at 50%. Quits in the wait zone are presented as solid black dots in (a,c) or as separate % curves in (b,d). % curves in (b,d) for enter and earn are plotted, with 100-% mirror curves for skip and quit intended to highlight cross over points that approximate threshold fits, particularly for later timepoints as animals discriminate tones in the offer zone but generally retain stable levels of willingness to wait in the wait zone. **e-h** Offer zone and wait zone thresholds plotted each day during the transition periods from reward-rich to reward-scarce environments (days 8-21) split across flavor rankings and split among the stepwise and gradual schedules and split into two separate 1-week windows reflecting the 1-5 s offer range (e) and 1-15 s offer range (f) for the stepwise schedule, with matched windows for the gradual schedule (g,h). Dashed green and blue lines represent the maximum possible thresholds on a given day for the offer zone and wait zone, respectively. Note in (h) STZ-gradual mice differ and deviate from VEH-gradual mice in two ways: (1) offer zone thresholds of STZ-gradual mice drift apart and are lower than VEH-gradual mice for less preferred flavors (*F*=9.491, *p*<0.0001) while (2) wait zone thresholds of STZ-gradual mice for most preferred flavors continue to scale with the changing max offer in the environment across days compared to VEH-gradual mice who peel off beginning on day 15 (*F*=3.797, *p*<0.0001), matching flat patterns of mice tested on the stepwise schedule in (f, treatment x rank x day: offer zone: *F*=0.381, *p*=0.767; wait zone: *F*=2.039, *p*=0.107). No effect of treatment on thresholds between days 8-14 of testing for stepwise (e, offer zone: *F*=0.371, *p*=0.774; wait zone: *F*=0.112, *p*=0.953) or gradual schedules (g, offer zone: *F*=1.091, *p*=0.353; wait zone: *F*=0.314, *p*=0.815). **i-p** Within-session meal consumption patterns collapsed across the entire 1-30 s epoch, split across 2.5 min time bins split by flavor rankings in (i-l) with schedules collapsed in (i,j) or separated in (k,l). Rewards earned in (i,k) or cumulative earns in (j,l). Data collapsed across flavors in (m,p). Shaded / error bars represent ±1 SEM. Shading in (e-h) represents 95% confidence interval of linear fit.

**Supplementary Fig. 4.**
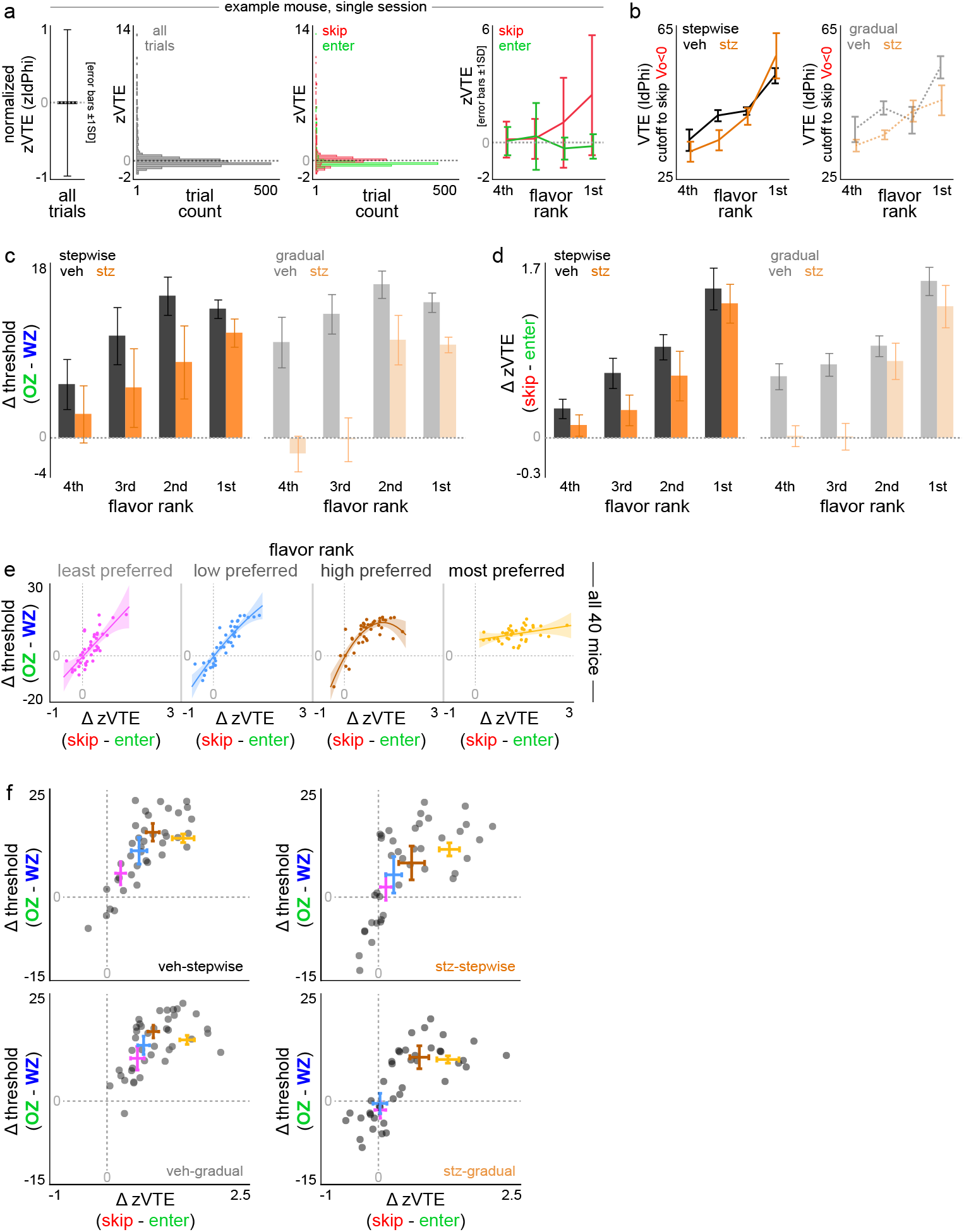
Expansion of choice conflict metrics on Restaurant Row. **a** Explanation of normalization of vicarious trial and error (VTE) behavior within animal. Z-score of VTE behavior calculated from all trials from a given mouse on a single day, regardless of flavor or choice outcome in the offer zone. Left panel illustrates that the average of this zVTE metric across all trials yields 0 plus or minus 1 standard deviation yielding values of plus or minus 1. Middle panels show the histogram distribution of zVTE metric across all trials in gray or trials subsequently split by enter (green) or skip (red) offer zone outcomes. Note the long tail for skip events. Average z-score values plotted split by flavor rankings and offer zone outcomes, with error bars here representing again standard deviation. Horizontal dashed gray lines represent z-score of 0. **b** The average amount of VTE displayed by mice before skipping negatively valued offers at least 50% of the time (Vo < 0, where offer value [Vo] = wait zone threshold – offer). This is termed VTE cutoff as previously reported (main effect of rank: *F*=26.980, *p*<0.0001, no effect of treatment: *F*=1.594, *p*=0.209 or schedule: *F*=0.595, *p*=0.442). **c** Difference score in offer zone minus wait zone thresholds, split by flavor ranking and schedule. **d** Difference score in zVTE skip minus enter choice outcomes, split by flavor ranking and schedule. **e** Scatter plot of difference scores for all 40 mice in (c) against (d) split by flavor ranking with quadratic curve fits showing a scaling of the ordinal rankings of conflict measures across flavor but with additional unexplained VTE differences not captured in threshold differences for more preferred flavors. **f** Data from (e) replotted but splitting groups of mice, with flavor ranking means and X-Y errors Colors in (e-f) are meant to help visually separate flavor rankings but do not actually reflect the specific flavors, as those are different from animal to animal. depicted. Data from (b-f) collapsed across the entire 1-30 s epoch. Shading in (e) represents 95% confidence interval of quadratic fit. Error bars in (b-d,f) represent ±1 SEM. Dots represent individual mice once in each panel of (e, within flavor) but each animal appears as 4 dots in (f, across each flavor superimposed).

**Supplementary Fig. 5.**
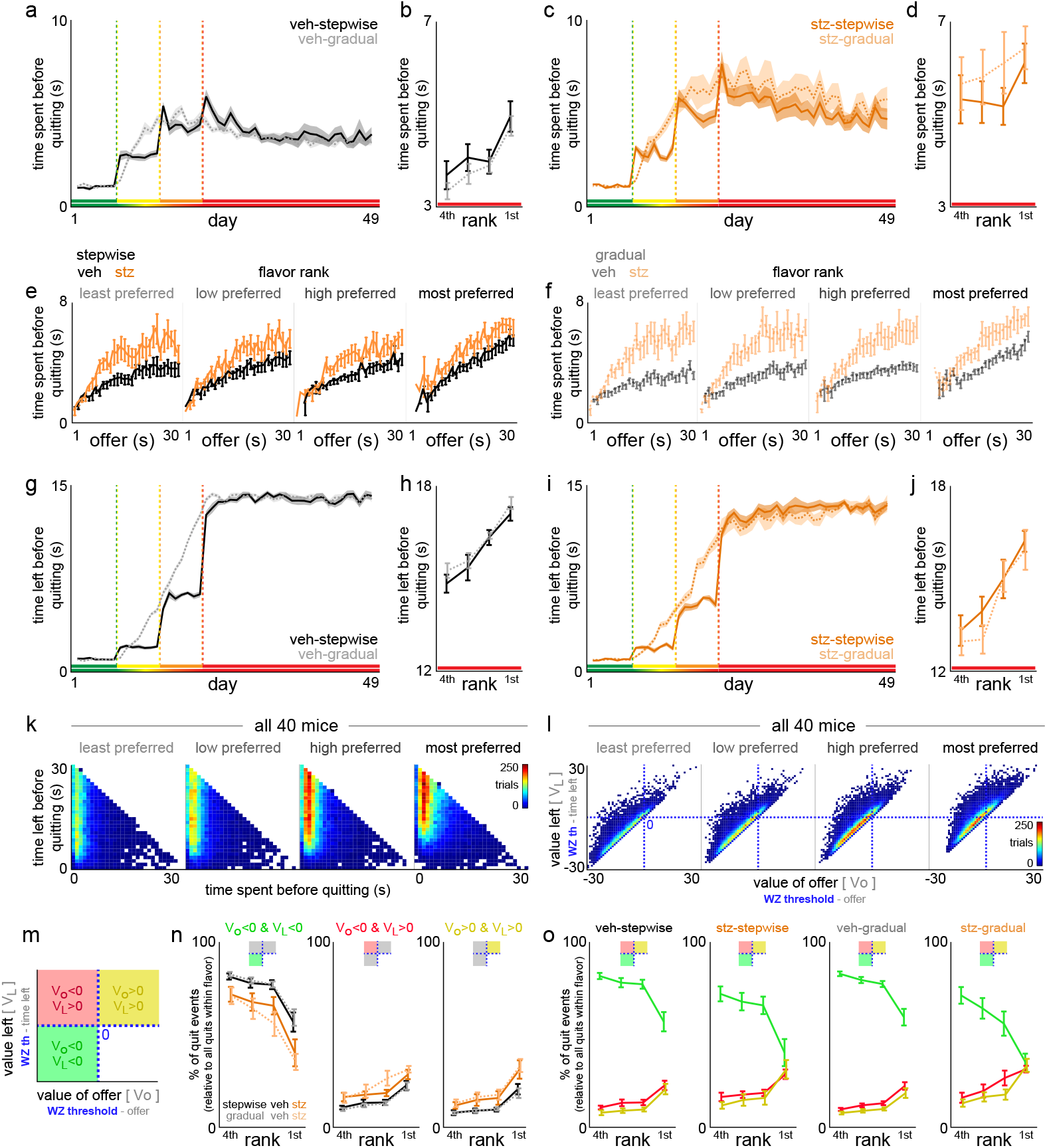
Expansion of quit metrics on Restaurant Row. **a-d** Latency to quit in the wait zone from countdown onset (enter decision) to the moment of quitting and exiting the wait zone mid-countdown (a,c) across the entire Restaurant Row paradigm or (b,d) collapsed across the 1-30 s epoch split by flavor rankings (main effect of treatment: *F*=17.546, *p*<0.0001; VEH rank: *F*=10.541, *p*<0.0001; STZ rank: *F*=0.993, *p*=0.401). **e-f** Data from (b,d) split by the original starting offer preceding each quit event. **g-j** Data from (a-d) but instead plotting how much time left was remaining in the countdown at the moment of quitting. **k** Pooled data across all mice from the 1-30 s epoch showing the distribution of trial counts of quit events derived from each unique [time spent, time left] pairs across all permutations derived from trials of various starting offers, split by flavor rankings. **l** Pooled data across all mice from the 1-30 s epoch split by flavor ranking showing all quit events categorized in value terms, both the value of the original starting offer (offer value [Vo] = wait zone threshold – offer) and the value of the amount of time left remaining in the countdown at the moment of quitting (value left [VL] = wait zone threshold – time left). Thus, this produces 4 quadrants, only three of which contain existing data. **m** Three categories of quit events based on offer value and value left terms. The green quadrant (Vo<0 & VL<0) represent the most efficient quit decisions from economically disadvantageous offers that in theory should have been skipped instead. Efficient quits describe quitting fast enough before it otherwise would have been more consistent with crossing one’s wait zone threshold to simply finish waiting instead. **n-o** Percentage of all quit events that derive from each quit category split by quit type in (n) or superimposed in (o) but split by groups of mice. Quit type x rank: *F*=49.958, *p*<0.0001. Note the interaction between quit type and treatment (*F*=8.645, *p*<0.001) but not schedule (*F*=0.044, *p*=0.957). These data do not capture the added change in quitting behavior that the sunk cost analysis captures (more dynamic changes due to the passage of time). Shaded / error bars represent ±1 SEM.

**Supplementary Fig. 6.**
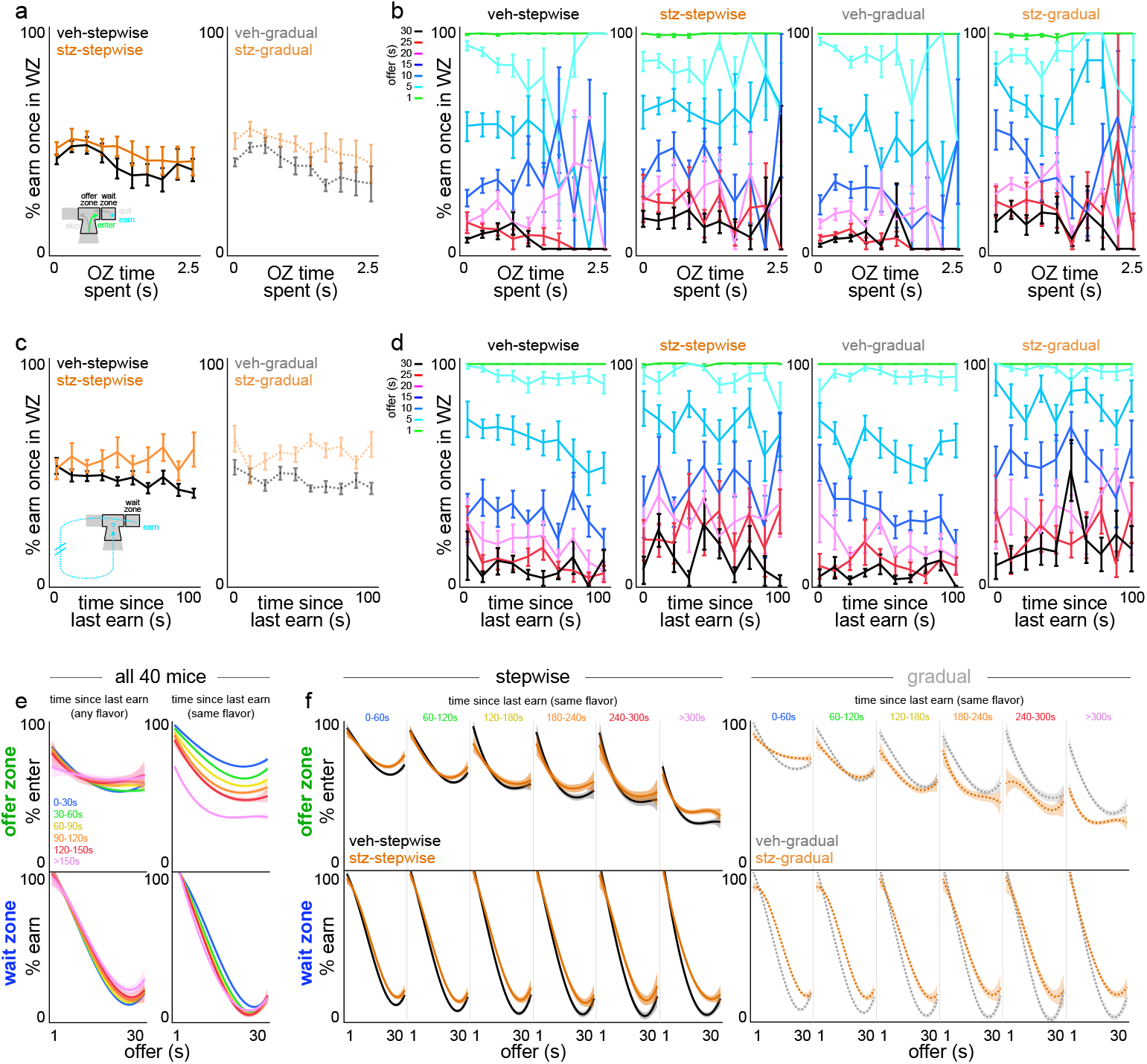
Valuing the passage of other forms of time on Restaurant Row. **a-b** Time spent in the offer zone before making an enter decision (e.g., fast vs. slow decisions in the offer zone) has no significant bearing on the likelihood of staying vs. quitting in the wait zone once entered either (a) overall or (b) segmented by what the starting offer of the delay was in color (relatively flat horizontal lines for each color that scale vertically reflecting the overall likelihood of quitting low-vs. high-delay offers, *F*=0.571, *p*=0.450). Fewer samples exist at longer OZ time spent, as animals are more likely to skip in the offer zone instead. **c-d** Time spent navigating around the maze since last reward was earned of any flavor (e.g., recently ate vs. it has been some time) has no significant bearing on the likelihood of staying vs. quitting in the wait zone once entered either (a) overall or (b) segmented by what the starting offer of the delay was in color upon arriving in the wait zone. While there is an effect of treatment x time (*F*=10.457, *p*<0.01), this effect does not significantly present across offers (*F*=0.292, *p*=0.589) nor interact with schedule (*F*=1.627, *p*=0.202). **e-f** Interestingly however, we find more complex value structure in this analysis by (1) whether or not the behavior in question is offer zone enter decisions vs. wait zone earn decisions and (2) whether or not the time metric is time elapsed since last earn of any flavor vs. time elapsed since last earn of the same flavor (calculated here first split by flavor ranking but then re-collapsed across flavors to show more simplified summary metrics). Data here is also plotted against cued offer costs along the x-axis to show psychometric curves of offer zone enter decisions or wait zone earn decisions as a function of cost. In (e), across all 40 mice, we only observe modulation of choice behavior in the offer zone when time elapsed considers time spent since last earn of a specific flavor. In (f), splitting this across schedules, with separate curves plotted for each window of time elapsed since last earn of the same flavor, we observe modulation of offer zone choice behavior differently between VEH- and STZ-treated mice only in those tested on the gradual schedule. Error bars represent ±1 SEM. Shading represents 95% confidence interval of cubic fit.

**Supplementary Fig. 7.**
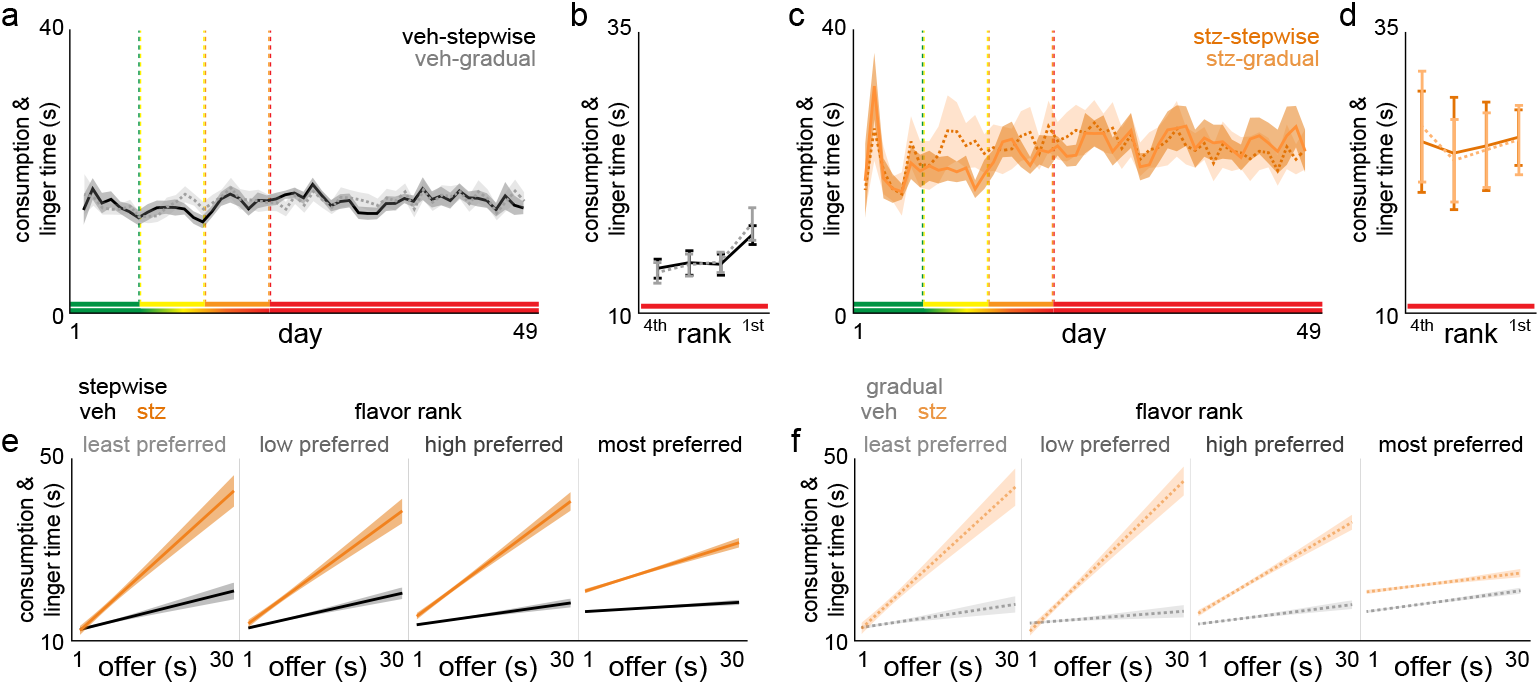
Post-earn behavior on Restaurant Row. After mice earn a reward, time was measured from pellet delivery onset until mice left the feeder site and exited the wait zone. This encompasses time spent consuming the reward as well as, as previously reported, additional time mice spent lingering at the reward site without engaging in any overt feeding behavior. **a-d** Consumption and linger time plotted (a,c) across the entire Restaurant Row paradigm or (b,d) split by flavor rankings collapsed across the 1-30 s epoch. **e-f** Data in (b,d) split by the original starting offer of the cued cost required to earn the reward on each trial, split by flavor rankings. We previously reported this behavior includes a component of a within-trial conditioned-place-preference-like behavior that carries some hedonic value or appraisal of the reward just consumed, as well as is sensitive to the amount time invested prior to earning the reward, even within flavor. Shaded / error bars represent ±1 SEM. Shading in (e-f) represents 95% confidence interval of linear fit.

**Supplementary Fig. 8.**
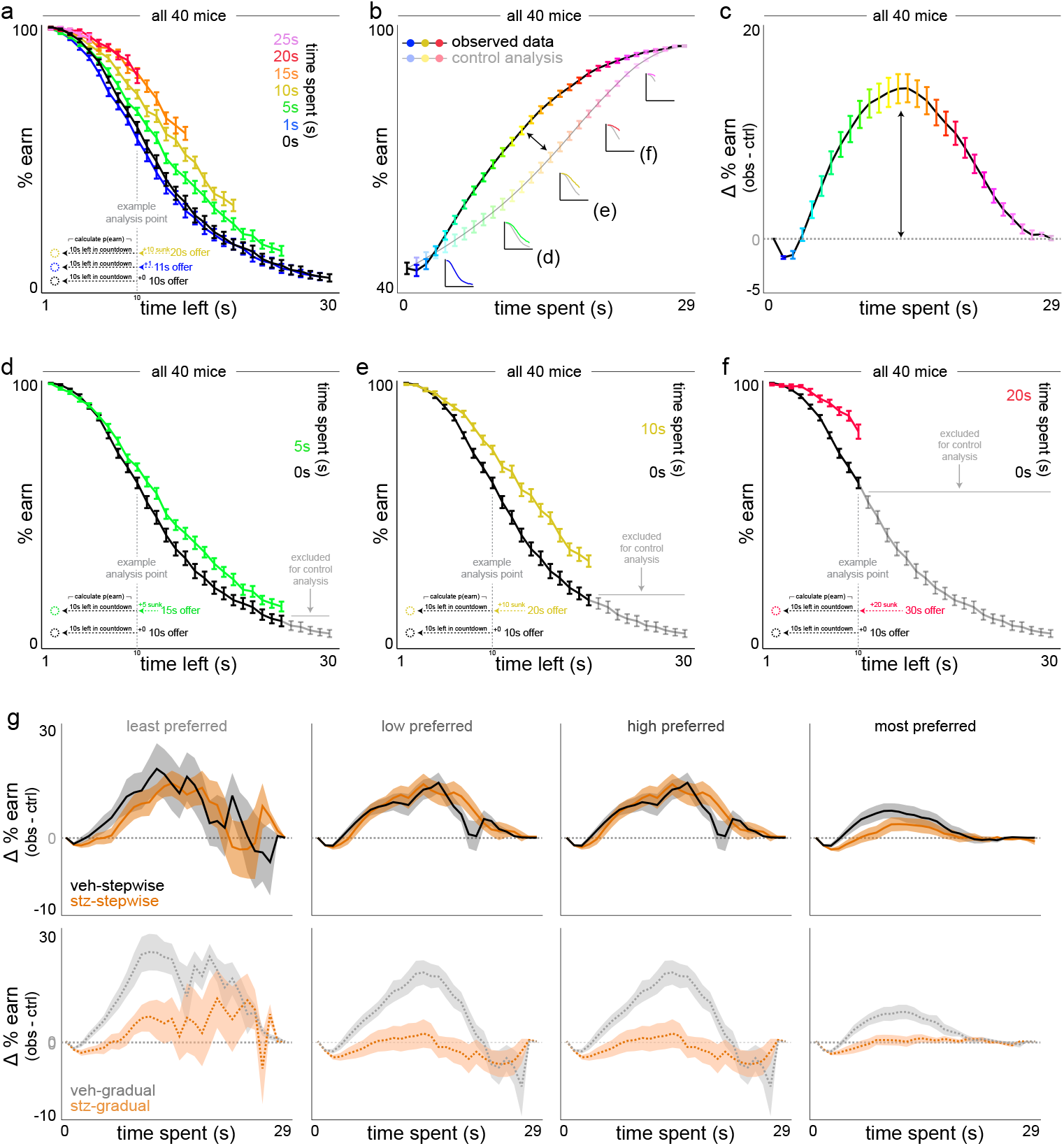
Visual explanation of sensitivity to sunk costs analysis in the wait zone and expanded data by flavor. **a-c** Sunk cost analysis of staying behavior in the wait zone, demonstrated using all mice. (a) The likelihood of staying in the wait zone and earning a reward (e.g., not quit) is plotted as a function of time left in the countdown along the x-axis and time already spent waiting orthogonally in color. Note the black 0 s time spent curve represents animals having just entered the wait zone from the offer zone. Inset vertical dashed gray line illustrates an example analysis point comparing three sunk cost conditions originating from different starting offers but matched at 10 s left. Data from (a) dimensioned reduced in (b) collapsing across time left, instead highlighting the grand mean of each time spent sunk cost condition (color and x-axis). Insets depict data from curves in (a) are collapsed into the observed (sunk condition) and control (0 s condition) lines. Difference between curves in (b) are plotted in (c) in order to summarize the envelope of the overall effect of time already spent on escalating the commitment of staying in the wait zone. Horizontal dashed line represents 0. **d-f** Redisplay data from (a) but for three sunk cost conditions indicated in the insets in (b, 5 s green in d, 10 s gold in e, 20 s red in f) with the 0 sunk cost condition (black curve) repeated in each panel. Because each colored sunk cost curve derives from trials where the starting offer cost is higher than the matched time left value of the 0 s cost condition, some data points do not exist and are missing from the sunk cost curve on the rightward end. Thus, when reducing dimensions, to control for the inflated summary calculations of % earn due to missing data alone, the control analysis iteratively excludes matched data from the black 0 s sunk curve (highlighted in gray) before collapsing data to yield the resultant control curve in (b). **g** Delta curves for VEH- and STZ-treated mice split by schedule and flavor rankings. Shaded / error bars represent ±1 SEM.

**Supplementary Fig. 9.**
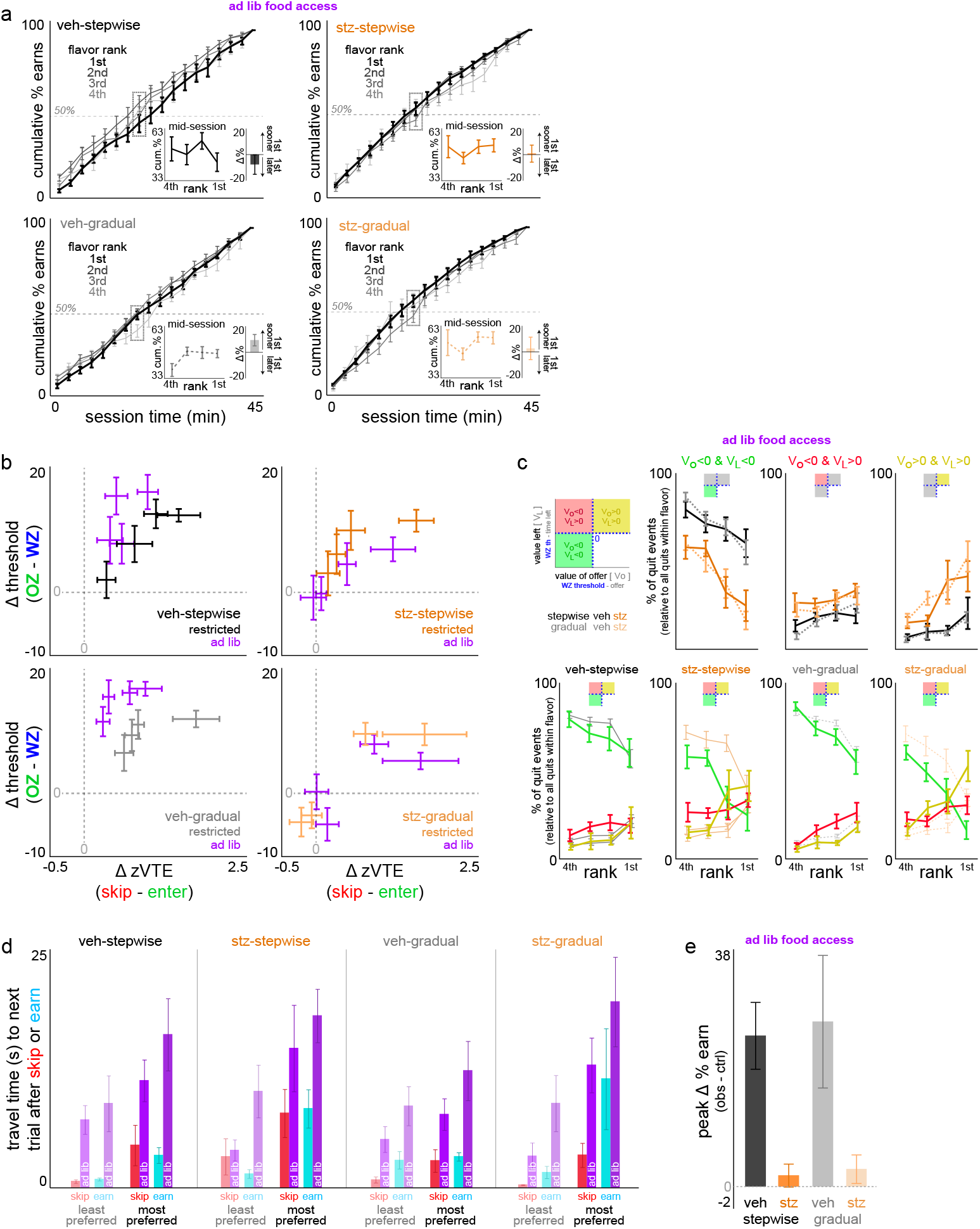
Expansion of additional Restaurant Row behavior while ad lib access to food was restored in the home cage. **a** Meal consumption patterns as in Fig. 4e. Note the overall gross collapsing of flavor patterns across all groups (sign test mid-session delta score against zero: VEH-stepwise: *t*=-1.020, *p*=0.335; VEH-gradual: *t*=2.140, *p*=0.061; STZ-stepwise: *t*=0.721, *p*=0.490; STZ-gradual: *t*=0.203, *p*=0.843) compared to restricted patterns as in Fig. 4e. **b** Difference scores in vicarious trial and error (VTE) choice conflict in the offer zone skip minus enter plotted against difference scores in offer zone minus wait zone threshold policies as in Supplementary Fig. 4f, comparing food restricted baselines (color of each group) to ad lib data (purple). Flavor rankings split and superimposed on top of one another as in Supplementary Fig. 4f (no flavor-specific visual indicators are labeled intentionally to highlight the gestalt of bidirectional shifts among VEH-vs. STZ-treated mice driven largely by changes only in wait zone thresholds and not offer zone thresholds (largely a y-axis shift), with relatively less changing along the x-axis in VTE (although with some decrease observable in VEH-treated mice), as depicted in Fig. 8d-f. **c** Categorization of quit types as in Supplementary Fig. 5m-o. Top shows data split by quit type, only depicting ad lib access data only. Bottom superimposes quit type separated by groups of mice. For visual reference, superimposed in the background in thinner lines in the color of each group of mice is the baseline restricted data while the ad lib data is plotted in the color of each quit type (green, gold, red) to appreciate no changes in VEH-treated mice while there are changes in STZ-treated mice as a result of ad lib access. Of note, STZ-gradual mice show the greatest change in distribution of quit types particularly for most preferred flavors. **d** Differences in travel time between restaurants in the hallways split by groups of mice, after exiting least (faint) vs. most (opaque) preferred restaurants, and immediately after just skipping (red) or earning and consuming a reward (cyan). Restricted baseline data in red/cyan while the corresponding ad lib data is adjacent in purple. Note the overall slowing in travel times due to ad lib access (*F*=22.657, *p*<0.0001). Also note the value structure in anticipatory travel times that interact with heading toward the next flavor after leaving least (faster) vs. most preferred restaurants (slower, *F*=49.402, *p*<0.0001), in spite of global differences in speed after skipping (faster) vs. having just eaten (slower, *F*=5.984, *p*<0.05). **e** Larger visual redisplay of the same data plotted as insets in Fig. 8g. Error bars represent ±1 SEM.

**Supplementary Fig. 10.**
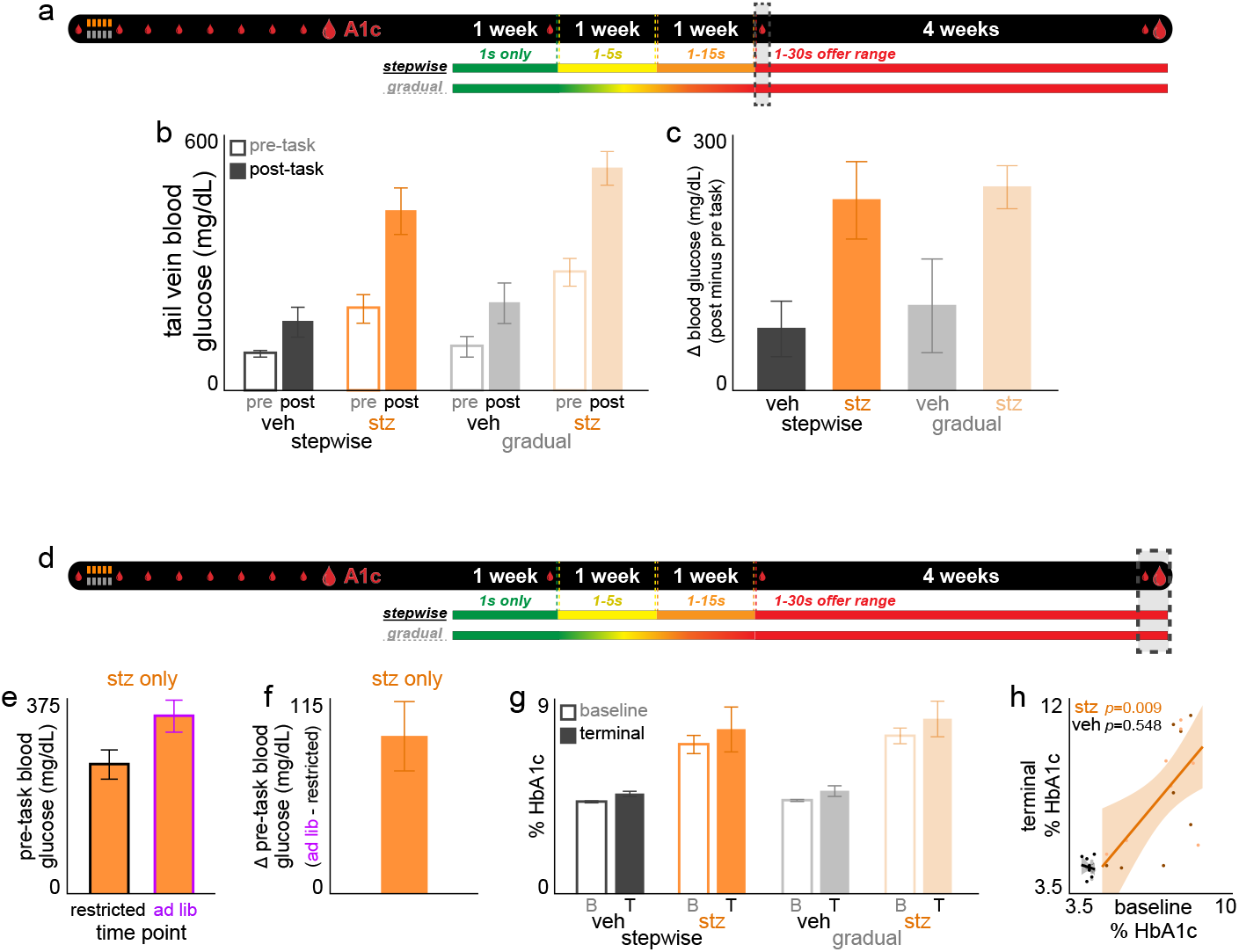
Additional timepoints of sampling blood glucose levels longitudinally across the entire Restaurant Row paradigm. **a** Timeline. Dashed box indicates time period (day 22) relevant for panels (b-c). This includes tail vein pre- and post-task samples of blood glucose levels (small droplet icon) for all 40 mice. **b** Pre- and post-task blood glucose levels. **c** Change in blood glucose levels post minus pre task. **d** Timeline. Dashed box indicates time period relevant for panels (e-h). This includes the final day of the experiment before sacrificing mice to extract terminal HbA1c levels (large droplet icon) as well as additional samples of tail vein pre-task only blood glucose levels obtained from STZ-mice only during the additional days of testing with ad lib access to regular chow after day 49 of the main experiment. **e** Pre-task blood glucose levels comparing ad lib access to restricted [using day 22] only in STZ-mice sampled at both timepoints. **f** Change in blood glucose from (e) ad lib minus restricted [pre-task only]. **g** HbA1c levels measured at baseline (before ever starting the entire Restaurant Row paradigm but after the initial 8-week STZ incubation [first large droplet in the timeline]) and terminally [second large droplet in the timeline]. **h** Data in (g) replotting time points against one another for all mice (darker dots visualize stepwise schedule). Error bars represent ±1 SEM. Shading represents 95% confidence interval of linear fit.

