## Supplementary Figures for "Diabetes alters neuroeconomically dissociable forms of mental accounting"

**Included:**
Supplementary Figures 1-10

Supplementary Figure 1

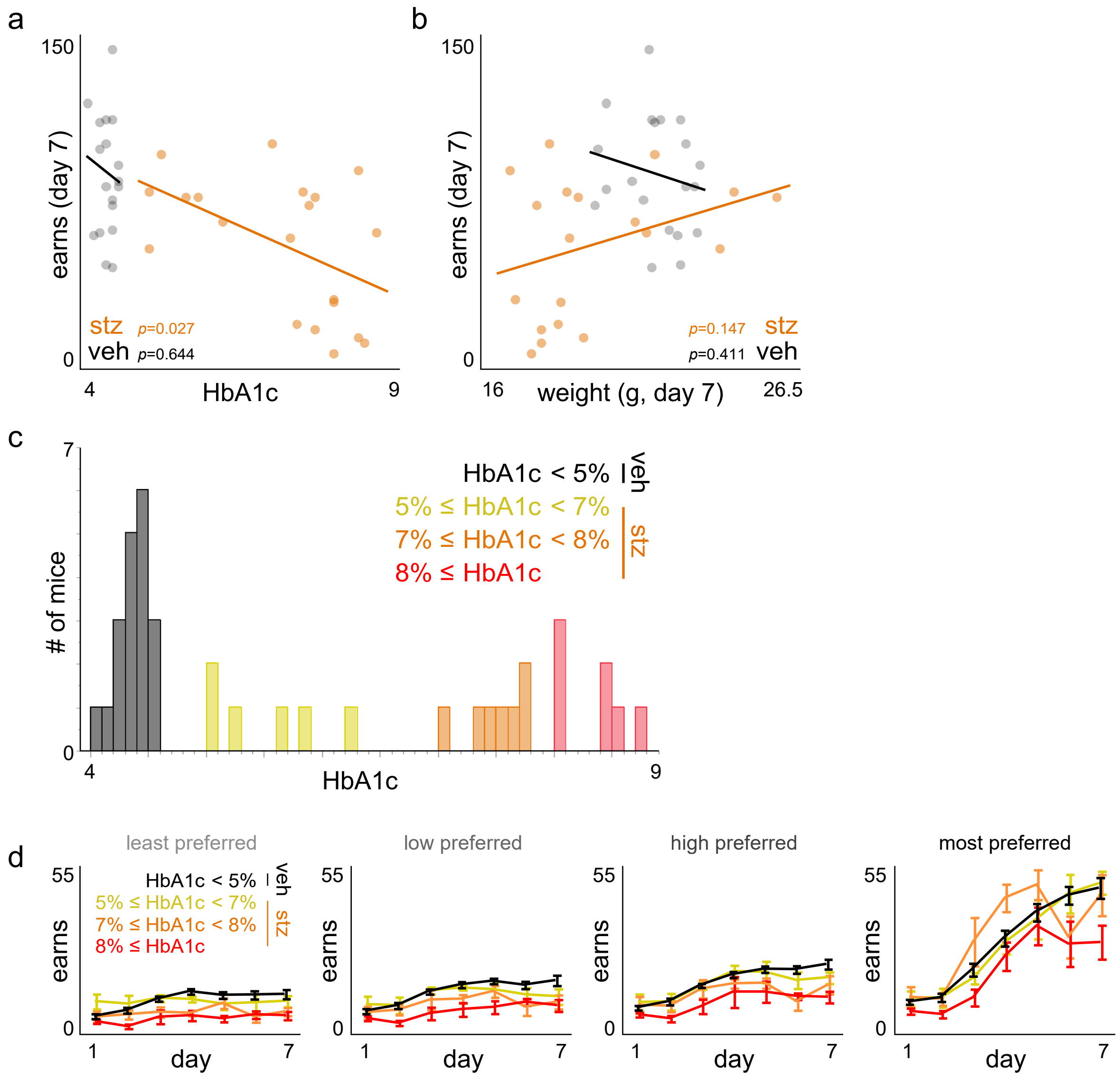

**Supplementary Fig. 1 | Analysis of HbA1c levels and behavior. a-b** Scatter plot of (a) HbA1c levels or (b) pre-task body weight measured on day 7 immediately before behavioral testing against total rewards earned on day 7. Trendline represents linear fit. **c** HbA1c levels split into evenly distributed sub-groups. **d** Visualization of rewards earned across days 1-7 split by flavor rankings in each HbA1c sub-group. Dots represent individual mice. Error bars represent ±1 SEM.

Supplementary Figure 2

**a**

| DAY | 1 | 2 | 3 | 4 | 5 | 6 | 7 | 8 | 9 | 10 | 11 | 12 | 13 | 14 | 15 | 16 | 17 | 18 | 19 | 20 | 21 | 22 | 23 | 24 | 25 | 26 | 27 | 28 | 29 | 30 | 31 | 32 | 33 | 34 | 35 | 36 | 37 | 38 | 39 | 40 | 41 | 42 | 43 | 44 | 45 | 46 | 47 | 48 | 49 |
|---|---|---|---|---|---|---|---|---|---|---|---|---|---|---|---|---|---|---|---|---|---|---|---|---|---|---|---|---|---|---|---|---|---|---|---|---|---|---|---|---|---|---|---|---|---|---|---|---|---|

**b** stepwise

| | | | | | | | | | | | | | | | | | | | | | | | | | | | | | | | | | | | | | | | | | | | | | | | | | |
|---|---|---|---|---|---|---|---|---|---|---|---|---|---|---|---|---|---|---|---|---|---|---|---|---|---|---|---|---|---|---|---|---|---|---|---|---|---|---|---|---|---|---|---|---|---|---|---|---|---|
| max offer |1|1|1|1|1|1|1|5|5|5|5|5|5|5|15|15|15|15|15|15|15|30|30|30|30|30|30|30|30|30|30|30|30|30|30|30|30|30|30|30|30|30|30|30|30|30|30|30|30|

**c**

| offer |1|2|3|4|5|6|7|8|9|10|11|12|13|14|15|16|17|18|19|20|21|22|23|24|25|26|27|28|29|30|31|32|33|34|35|36|37|38|39|40|41|42|43|44|45|46|47|48|49|
|---|---|---|---|---|---|---|---|---|---|---|---|---|---|---|---|---|---|---|---|---|---|---|---|---|---|---|---|---|---|---|---|---|---|---|---|---|---|---|---|---|---|---|---|---|---|---|---|---|---|
| 1s |1|2|3|4|5|6|7|8|9|10|11|12|13|14|15|16|17|18|19|20|21|22|23|24|25|26|27|28|29|30|31|32|33|34|35|36|37|38|39|40|41|42|43|44|45|46|47|48|49|
| 2s |0|0|0|0|0|0|0|1|2|3|4|5|6|7|8|9|10|11|12|13|14|15|16|17|18|19|20|21|22|23|24|25|26|27|28|29|30|31|32|33|34|35|36|37|38|39|40|41|42|
| 3s |0|0|0|0|0|0|0|1|2|3|4|5|6|7|8|9|10|11|12|13|14|15|16|17|18|19|20|21|22|23|24|25|26|27|28|29|30|31|32|33|34|35|36|37|38|39|40|41|42|
| 4s |0|0|0|0|0|0|0|1|2|3|4|5|6|7|8|9|10|11|12|13|14|15|16|17|18|19|20|21|22|23|24|25|26|27|28|29|30|31|32|33|34|35|36|37|38|39|40|41|42|
| 5s |0|0|0|0|0|0|0|1|2|3|4|5|6|7|8|9|10|11|12|13|14|15|16|17|18|19|20|21|22|23|24|25|26|27|28|29|30|31|32|33|34|35|36|37|38|39|40|41|42|
| 6s |0|0|0|0|0|0|0|0|0|0|0|0|0|0|1|2|3|4|5|6|7|8|9|10|11|12|13|14|15|16|17|18|19|20|21|22|23|24|25|26|27|28|29|30|31|32|33|34|35|
| 7s |0|0|0|0|0|0|0|0|0|0|0|0|0|0|1|2|3|4|5|6|7|8|9|10|11|12|13|14|15|16|17|18|19|20|21|22|23|24|25|26|27|28|29|30|31|32|33|34|35|
| 8s |0|0|0|0|0|0|0|0|0|0|0|0|0|0|1|2|3|4|5|6|7|8|9|10|11|12|13|14|15|16|17|18|19|20|21|22|23|24|25|26|27|28|29|30|31|32|33|34|35|
| 9s |0|0|0|0|0|0|0|0|0|0|0|0|0|0|1|2|3|4|5|6|7|8|9|10|11|12|13|14|15|16|17|18|19|20|21|22|23|24|25|26|27|28|29|30|31|32|33|34|35|
| 10s |0|0|0|0|0|0|0|0|0|0|0|0|0|0|1|2|3|4|5|6|7|8|9|10|11|12|13|14|15|16|17|18|19|20|21|22|23|24|25|26|27|28|29|30|31|32|33|34|35|
| 11s |0|0|0|0|0|0|0|0|0|0|0|0|0|0|1|2|3|4|5|6|7|8|9|10|11|12|13|14|15|16|17|18|19|20|21|22|23|24|25|26|27|28|29|30|31|32|33|34|35|
| 12s |0|0|0|0|0|0|0|0|0|0|0|0|0|0|1|2|3|4|5|6|7|8|9|10|11|12|13|14|15|16|17|18|19|20|21|22|23|24|25|26|27|28|29|30|31|32|33|34|35|
| 13s |0|0|0|0|0|0|0|0|0|0|0|0|0|0|1|2|3|4|5|6|7|8|9|10|11|12|13|14|15|16|17|18|19|20|21|22|23|24|25|26|27|28|29|30|31|32|33|34|35|
| 14s |0|0|0|0|0|0|0|0|0|0|0|0|0|0|1|2|3|4|5|6|7|8|9|10|11|12|13|14|15|16|17|18|19|20|21|22|23|24|25|26|27|28|29|30|31|32|33|34|35|
| 15s |0|0|0|0|0|0|0|0|0|0|0|0|0|0|1|2|3|4|5|6|7|8|9|10|11|12|13|14|15|16|17|18|19|20|21|22|23|24|25|26|27|28|29|30|31|32|33|34|35|
| 16s |0|0|0|0|0|0|0|0|0|0|0|0|0|0|0|0|0|0|0|0|0|1|2|3|4|5|6|7|8|9|10|11|12|13|14|15|16|17|18|19|20|21|22|23|24|25|26|27|28|
| 17s |0|0|0|0|0|0|0|0|0|0|0|0|0|0|0|0|0|0|0|0|0|1|2|3|4|5|6|7|8|9|10|11|12|13|14|15|16|17|18|19|20|21|22|23|24|25|26|27|28|
| 18s |0|0|0|0|0|0|0|0|0|0|0|0|0|0|0|0|0|0|0|0|0|1|2|3|4|5|6|7|8|9|10|11|12|13|14|15|16|17|18|19|20|21|22|23|24|25|26|27|28|
| 19s |0|0|0|0|0|0|0|0|0|0|0|0|0|0|0|0|0|0|0|0|0|1|2|3|4|5|6|7|8|9|10|11|12|13|14|15|16|17|18|19|20|21|22|23|24|25|26|27|28|
| 20s |0|0|0|0|0|0|0|0|0|0|0|0|0|0|0|0|0|0|0|0|0|1|2|3|4|5|6|7|8|9|10|11|12|13|14|15|16|17|18|19|20|21|22|23|24|25|26|27|28|
| 21s |0|0|0|0|0|0|0|0|0|0|0|0|0|0|0|0|0|0|0|0|0|1|2|3|4|5|6|7|8|9|10|11|12|13|14|15|16|17|18|19|20|21|22|23|24|25|26|27|28|
| 22s |0|0|0|0|0|0|0|0|0|0|0|0|0|0|0|0|0|0|0|0|0|1|2|3|4|5|6|7|8|9|10|11|12|13|14|15|16|17|18|19|20|21|22|23|24|25|26|27|28|
| 23s |0|0|0|0|0|0|0|0|0|0|0|0|0|0|0|0|0|0|0|0|0|1|2|3|4|5|6|7|8|9|10|11|12|13|14|15|16|17|18|19|20|21|22|23|24|25|26|27|28|
| 24s |0|0|0|0|0|0|0|0|0|0|0|0|0|0|0|0|0|0|0|0|0|1|2|3|4|5|6|7|8|9|10|11|12|13|14|15|16|17|18|19|20|21|22|23|24|25|26|27|28|
| 25s |0|0|0|0|0|0|0|0|0|0|0|0|0|0|0|0|0|0|0|0|0|1|2|3|4|5|6|7|8|9|10|11|12|13|14|15|16|17|18|19|20|21|22|23|24|25|26|27|28|
| 26s |0|0|0|0|0|0|0|0|0|0|0|0|0|0|0|0|0|0|0|0|0|1|2|3|4|5|6|7|8|9|10|11|12|13|14|15|16|17|18|19|20|21|22|23|24|25|26|27|28|
| 27s |0|0|0|0|0|0|0|0|0|0|0|0|0|0|0|0|0|0|0|0|0|1|2|3|4|5|6|7|8|9|10|11|12|13|14|15|16|17|18|19|20|21|22|23|24|25|26|27|28|
| 28s |0|0|0|0|0|0|0|0|0|0|0|0|0|0|0|0|0|0|0|0|0|1|2|3|4|5|6|7|8|9|10|11|12|13|14|15|16|17|18|19|20|21|22|23|24|25|26|27|28|
| 29s |0|0|0|0|0|0|0|0|0|0|0|0|0|0|0|0|0|0|0|0|0|1|2|3|4|5|6|7|8|9|10|11|12|13|14|15|16|17|18|19|20|21|22|23|24|25|26|27|28|
| 30s |0|0|0|0|0|0|0|0|0|0|0|0|0|0|0|0|0|0|0|0|0|1|2|3|4|5|6|7|8|9|10|11|12|13|14|15|16|17|18|19|20|21|22|23|24|25|26|27|28|

**d** gradual

| | | | | | | | | | | | | | | | | | | | | | | | | | | | | | | | | | | | | | | | | | | | | | | | | | |
|---|---|---|---|---|---|---|---|---|---|---|---|---|---|---|---|---|---|---|---|---|---|---|---|---|---|---|---|---|---|---|---|---|---|---|---|---|---|---|---|---|---|---|---|---|---|---|---|---|---|
| max offer |1|1|1|1|1|1|1|2|4|6|8|10|12|14|16|18|20|22|24|26|28|30|30|30|30|30|30|30|30|30|30|30|30|30|30|30|30|30|30|30|30|30|30|30|30|30|30|30|30|

**e**

| offer |1|2|3|4|5|6|7|8|9|10|11|12|13|14|15|16|17|18|19|20|21|22|23|24|25|26|27|28|29|30|31|32|33|34|35|36|37|38|39|40|41|42|43|44|45|46|47|48|49|
|---|---|---|---|---|---|---|---|---|---|---|---|---|---|---|---|---|---|---|---|---|---|---|---|---|---|---|---|---|---|---|---|---|---|---|---|---|---|---|---|---|---|---|---|---|---|---|---|---|---|
| 1s |1|2|3|4|5|6|7|8|9|10|11|12|13|14|15|16|17|18|19|20|21|22|23|24|25|26|27|28|29|30|31|32|33|34|35|36|37|38|39|40|41|42|43|44|45|46|47|48|49|
| 2s |0|0|0|0|0|0|0|1|2|3|4|5|6|7|8|9|10|11|12|13|14|15|16|17|18|19|20|21|22|23|24|25|26|27|28|29|30|31|32|33|34|35|36|37|38|39|40|41|42|
| 3s |0|0|0|0|0|0|0|0|1|2|3|4|5|6|7|8|9|10|11|12|13|14|15|16|17|18|19|20|21|22|23|24|25|26|27|28|29|30|31|32|33|34|35|36|37|38|39|40|41|
| 4s |0|0|0|0|0|0|0|0|1|2|3|4|5|6|7|8|9|10|11|12|13|14|15|16|17|18|19|20|21|22|23|24|25|26|27|28|29|30|31|32|33|34|35|36|37|38|39|40|41|
| 5s |0|0|0|0|0|0|0|0|0|1|2|3|4|5|6|7|8|9|10|11|12|13|14|15|16|17|18|19|20|21|22|23|24|25|26|27|28|29|30|31|32|33|34|35|36|37|38|39|40|
| 6s |0|0|0|0|0|0|0|0|0|1|2|3|4|5|6|7|8|9|10|11|12|13|14|15|16|17|18|19|20|21|22|23|24|25|26|27|28|29|30|31|32|33|34|35|36|37|38|39|40|
| 7s |0|0|0|0|0|0|0|0|0|0|1|2|3|4|5|6|7|8|9|10|11|12|13|14|15|16|17|18|19|20|21|22|23|24|25|26|27|28|29|30|31|32|33|34|35|36|37|38|39|
| 8s |0|0|0|0|0|0|0|0|0|0|1|2|3|4|5|6|7|8|9|10|11|12|13|14|15|16|17|18|19|20|21|22|23|24|25|26|27|28|29|30|31|32|33|34|35|36|37|38|39|
| 9s |0|0|0|0|0|0|0|0|0|0|0|1|2|3|4|5|6|7|8|9|10|11|12|13|14|15|16|17|18|19|20|21|22|23|24|25|26|27|28|29|30|31|32|33|34|35|36|37|38|
| 10s |0|0|0|0|0|0|0|0|0|0|0|1|2|3|4|5|6|7|8|9|10|11|12|13|14|15|16|17|18|19|20|21|22|23|24|25|26|27|28|29|30|31|32|33|34|35|36|37|38|
| 11s |0|0|0|0|0|0|0|0|0|0|0|0|1|2|3|4|5|6|7|8|9|10|11|12|13|14|15|16|17|18|19|20|21|22|23|24|25|26|27|28|29|30|31|32|33|34|35|36|37|
| 12s |0|0|0|0|0|0|0|0|0|0|0|0|1|2|3|4|5|6|7|8|9|10|11|12|13|14|15|16|17|18|19|20|21|22|23|24|25|26|27|28|29|30|31|32|33|34|35|36|37|
| 13s |0|0|0|0|0|0|0|0|0|0|0|0|0|1|2|3|4|5|6|7|8|9|10|11|12|13|14|15|16|17|18|19|20|21|22|23|24|25|26|27|28|29|30|31|32|33|34|35|36|
| 14s |0|0|0|0|0|0|0|0|0|0|0|0|0|1|2|3|4|5|6|7|8|9|10|11|12|13|14|15|16|17|18|19|20|21|22|23|24|25|26|27|28|29|30|31|32|33|34|35|36|
| 15s |0|0|0|0|0|0|0|0|0|0|0|0|0|0|1|2|3|4|5|6|7|8|9|10|11|12|13|14|15|16|17|18|19|20|21|22|23|24|25|26|27|28|29|30|31|32|33|34|35|
| 16s |0|0|0|0|0|0|0|0|0|0|0|0|0|0|1|2|3|4|5|6|7|8|9|10|11|12|13|14|15|16|17|18|19|20|21|22|23|24|25|26|27|28|29|30|31|32|33|34|35|
| 17s |0|0|0|0|0|0|0|0|0|0|0|0|0|0|0|1|2|3|4|5|6|7|8|9|10|11|12|13|14|15|16|17|18|19|20|21|22|23|24|25|26|27|28|29|30|31|32|33|34|
| 18s |0|0|0|0|0|0|0|0|0|0|0|0|0|0|0|1|2|3|4|5|6|7|8|9|10|11|12|13|14|15|16|17|18|19|20|21|22|23|24|25|26|27|28|29|30|31|32|33|34|
| 19s |0|0|0|0|0|0|0|0|0|0|0|0|0|0|0|0|1|2|3|4|5|6|7|8|9|10|11|12|13|14|15|16|17|18|19|20|21|22|23|24|25|26|27|28|29|30|31|32|33|
| 20s |0|0|0|0|0|0|0|0|0|0|0|0|0|0|0|0|1|2|3|4|5|6|7|8|9|10|11|12|13|14|15|16|17|18|19|20|21|22|23|24|25|26|27|28|29|30|31|32|33|
| 21s |0|0|0|0|0|0|0|0|0|0|0|0|0|0|0|0|0|1|2|3|4|5|6|7|8|9|10|11|12|13|14|15|16|17|18|19|20|21|22|23|24|25|26|27|28|29|30|31|32|
| 22s |0|0|0|0|0|0|0|0|0|0|0|0|0|0|0|0|0|1|2|3|4|5|6|7|8|9|10|11|12|13|14|15|16|17|18|19|20|21|22|23|24|25|26|27|28|29|30|31|32|
| 23s |0|0|0|0|0|0|0|0|0|0|0|0|0|0|0|0|0|0|1|2|3|4|5|6|7|8|9|10|11|12|13|14|15|16|17|18|19|20|21|22|23|24|25|26|27|28|29|30|31|
| 24s |0|0|0|0|0|0|0|0|0|0|0|0|0|0|0|0|0|0|1|2|3|4|5|6|7|8|9|10|11|12|13|14|15|16|17|18|19|20|21|22|23|24|25|26|27|28|29|30|31|
| 25s |0|0|0|0|0|0|0|0|0|0|0|0|0|0|0|0|0|0|0|1|2|3|4|5|6|7|8|9|10|11|12|13|14|15|16|17|18|19|20|21|22|23|24|25|26|27|28|29|30|
| 26s |0|0|0|0|0|0|0|0|0|0|0|0|0|0|0|0|0|0|0|1|2|3|4|5|6|7|8|9|10|11|12|13|14|15|16|17|18|19|20|21|22|23|24|25|26|27|28|29|30|
| 27s |0|0|0|0|0|0|0|0|0|0|0|0|0|0|0|0|0|0|0|0|1|2|3|4|5|6|7|8|9|10|11|12|13|14|15|16|17|18|19|20|21|22|23|24|25|26|27|28|29|
| 28s |0|0|0|0|0|0|0|0|0|0|0|0|0|0|0|0|0|0|0|0|1|2|3|4|5|6|7|8|9|10|11|12|13|14|15|16|17|18|19|20|21|22|23|24|25|26|27|28|29|
| 29s |0|0|0|0|0|0|0|0|0|0|0|0|0|0|0|0|0|0|0|0|0|1|2|3|4|5|6|7|8|9|10|11|12|13|14|15|16|17|18|19|20|21|22|23|24|25|26|27|28|
| 30s |0|0|0|0|0|0|0|0|0|0|0|0|0|0|0|0|0|0|0|0|0|1|2|3|4|5|6|7|8|9|10|11|12|13|14|15|16|17|18|19|20|21|22|23|24|25|26|27|28|

**f**

| offer | stepwise | gradual | difference |
|---|---|---|---|
| 1s | 49 | 49 | 0 |
| 2s | 42 | 42 | 0 |
| 3s | 42 | 41 | -1 |
| 4s | 42 | 41 | -1 |
| 5s | 42 | 40 | -2 |
| 6s | 35 | 40 | 5 |
| 7s | 35 | 39 | 4 |
| 8s | 35 | 39 | 4 |
| 9s | 35 | 38 | 3 |
| 10s | 35 | 38 | 3 |
| 11s | 35 | 37 | 2 |
| 12s | 35 | 37 | 2 |
| 13s | 35 | 36 | 1 |
| 14s | 35 | 36 | 1 |
| 15s | 35 | 35 | 0 |
| 16s | 28 | 35 | 7 |
| 17s | 28 | 34 | 6 |
| 18s | 28 | 34 | 6 |
| 19s | 28 | 33 | 5 |
| 20s | 28 | 33 | 5 |
| 21s | 28 | 32 | 4 |
| 22s | 28 | 32 | 4 |
| 23s | 28 | 31 | 3 |
| 24s | 28 | 31 | 3 |
| 25s | 28 | 30 | 2 |
| 26s | 28 | 30 | 2 |
| 27s | 28 | 29 | 1 |
| 28s | 28 | 29 | 1 |
| 29s | 28 | 28 | 0 |
| 30s | 28 | 28 | 0 |
| days w/ 1s vs 15s | 7 | 7 | 0 |
| days w/ 1s vs 30s | 21 | 21 | 0 |

**Supplementary Fig. 2 | Visualization of experimentally manipulated Restaurant Row testing schedules. a** Timeline of the entire Restaurant Row paradigm, days 1 to 49 enumerated. Red marking on day 22 indicates the first day of the transition into the 1-30 s reward-scarce epoch, matched for both schedules. **b** Stepwise schedule timeline. Max offer on each day enumerated and color coded. **c** Stepwise schedule visualization of cumulative number of days animals would have had experience with each offer length, enumerated and color-coded scale ranging from 0 days of experience (green) to 49 days of experience (red). Staircase dashed line represents transition points. **d** Gradual schedule timeline. Max offer on each day enumerated and color coded. **e** Gradual schedule visualization of cumulative number of days animals would have had experience with each offer length, enumerated and color-coded scale ranging from 0 days of experience (green) to 49 days of experience (red). Diagonal dashed line represents gradual transition points. **f** Comparison of metrics between stepwise and gradual schedules. Columns 1 and 2 enumerate total number of days of experience with each offer length. Column 3 represents the difference in total number of days of experience with each offer length between stepwise minus gradual schedules (cool color-code). Bottom two rows of table indicate the difference between number of days of experience with 1 s offers vs 15 s offers or 1 s offers vs 30 s offers between the stepwise and gradual schedules, which are matched. This experimental design optimally controls for as many differences between schedules while varying rate of environmental change.

### Supplementary Figure 3

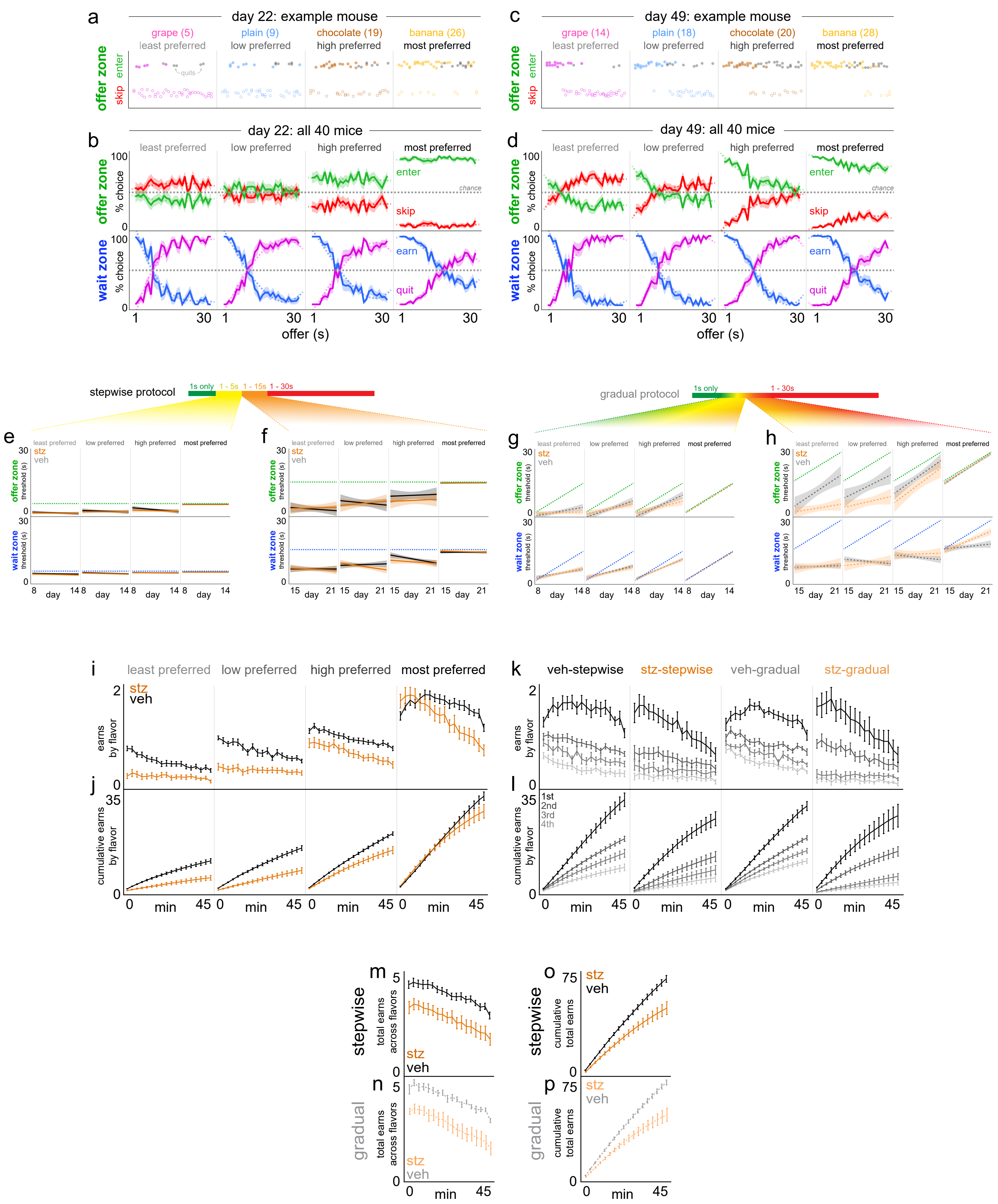

**Supplementary Fig. 3 | Expansion of choice behavior metrics on Restaurant Row. a-d** Example behavior from a single mouse on (a) day 22 and (c) day 49 as well as aggregate example summary choice data from all 40 mice on (b) day 22 and (d) day 49. Data depict offer zone choice outcomes on the y-axis (enter vs. skip) as a function of cued offer cost (tone pitch) on the x-axis either as individual choices (dots in a,c) or as summary % choice (curves in b,d). Horizontal dashed gray line in (b,d) indicates chance at 50%. Quits in the wait zone are presented as solid black dots in (a,c) or as separate % curves in (b,d). % curves in (b,d) for enter and earn are plotted, with 100-% mirror curves for skip and quit intended to highlight cross over points that approximate threshold fits, particularly for later timepoints as animals discriminate tones in the offer zone but generally retain stable levels of willingness to wait in the wait zone. **e-h** Offer zone and wait zone thresholds plotted each day during the transition periods from reward-rich to reward-scarce environments (days 8-21) split across flavor rankings and split among the stepwise and gradual schedules and split into two separate 1-week windows reflecting the 1-5 s offer range (e) and 1-15 s offer range (f) for the stepwise schedule, with matched windows for the gradual schedule (g,h). Dashed green and blue lines represent the maximum possible thresholds on a given day for the offer zone and wait zone, respectively. Note in (h) STZ-gradual mice differ and deviate from VEH-gradual mice in two ways: (1) offer zone thresholds of STZ-gradual mice drift apart and are lower than VEH-gradual mice for less preferred flavors ($F$=9.491, $p$<0.0001) while (2) wait zone thresholds of STZ-gradual mice for most preferred flavors continue to scale with the changing max offer in the environment across days compared to VEH-gradual mice who peel off beginning on day 15 ($F$=3.797, $p$<0.0001), matching flat patterns of mice tested on the stepwise schedule in (f, treatment x rank x day: offer zone: $F$=0.381, $p$=0.767; wait zone: $F$=2.039, $p$=0.107). No effect of treatment on thresholds between days 8-14 of testing for stepwise (e, offer zone: $F$=0.371, $p$=0.774; wait zone: $F$=0.112, $p$=0.953) or gradual schedules (g, offer zone: $F$=1.091, $p$=0.353; wait zone: $F$=0.314, $p$=0.815). **i-p** Within-session meal consumption patterns collapsed across the entire 1-30 s epoch, split across 2.5 min time bins split by flavor rankings in (i-l) with schedules collapsed in (i,j) or separated in (k,l). Rewards earned in (i,k) or cumulative earns in (j,l). Data collapsed across flavors in (m,p). Shaded / error bars represent ±1 SEM. Shading in (e-h) represents 95% confidence interval of linear fit.

#### Supplementary Figure 4

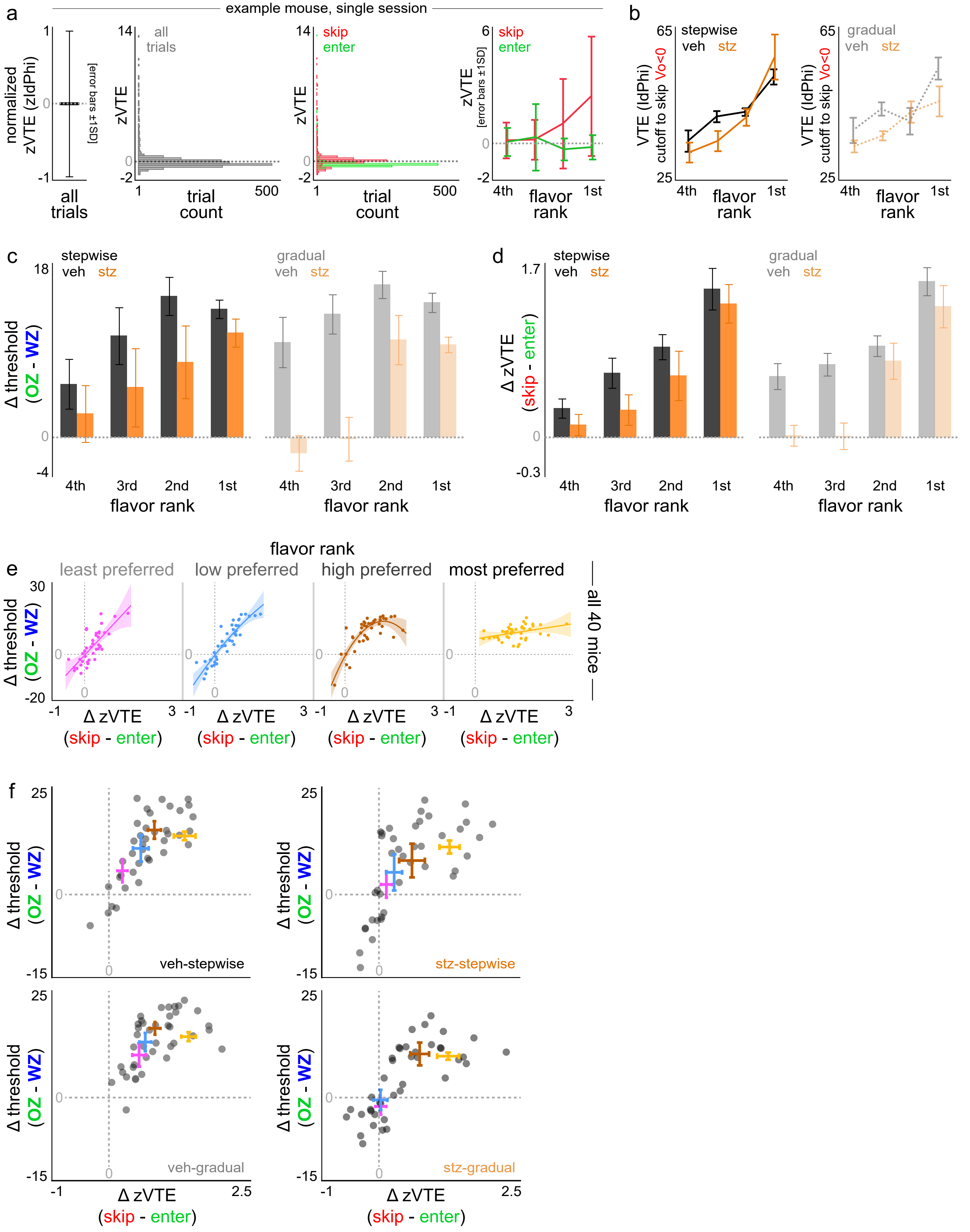

**Supplementary Fig. 4 | Expansion of choice conflict metrics on Restaurant Row. a** Explanation of normalization of vicarious trial and error (VTE) behavior within animal. Z-score of VTE behavior calculated from all trials from a given mouse on a single day, regardless of flavor or choice outcome in the offer zone. Left panel illustrates that the average of this zVTE metric across all trials yields 0 plus or minus 1 standard deviation yielding values of plus or minus 1. Middle panels show the histogram distribution of zVTE metric across all trials in gray or trials subsequently split by enter (green) or skip (red) offer zone outcomes. Note the long tail for skip events. Average z-score values plotted split by flavor rankings and offer zone outcomes, with error bars here representing again standard deviation. Horizontal dashed gray lines represent z-score of 0. **b** The average amount of VTE displayed by mice before skipping negatively valued offers at least 50% of the time (Vo < 0, where offer value [Vo] = wait zone threshold – offer). This is termed VTE cutoff as previously reported (main effect of rank: $F$=26.980, $p$<0.0001, no effect of treatment: $F$=1.594, $p$=0.209 or schedule: $F$=0.595, $p$=0.442). **c** Difference score in offer zone minus wait zone thresholds, split by flavor ranking and schedule. **d** Difference score in zVTE skip minus enter choice outcomes, split by flavor ranking and schedule. **e** Scatter plot of difference scores for all 40 mice in (c) against (d) split by flavor ranking with quadratic curve fits showing a scaling of the ordinal rankings of conflict measures across flavor but with additional unexplained VTE differences not captured in threshold differences for more preferred flavors. **f** Data from (e) replotted but splitting groups of mice, with flavor ranking means and X-Y errors Colors in (e-f) are meant to help visually separate flavor rankings but do not actually reflect the specific flavors, as those are different from animal to animal. depicted. Data from (b-f) collapsed across the entire 1-30 s epoch. Shading in (e) represents 95% confidence interval of quadratic fit. Error bars in (b-d,f) represent ±1 SEM. Dots represent individual mice once in each panel of (e, within flavor) but each animal appears as 4 dots in (f, across each flavor superimposed).

### Supplementary Figure 5

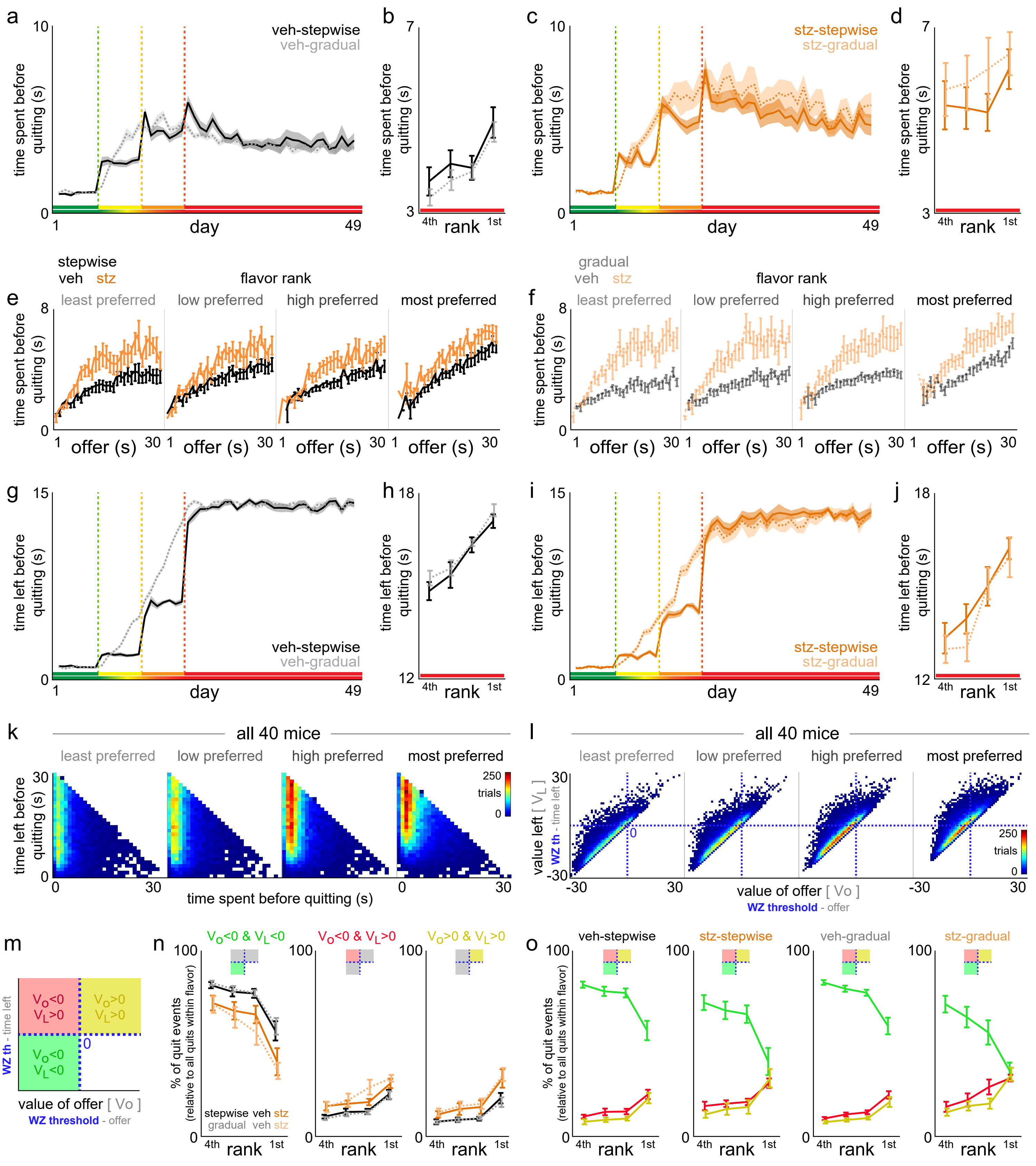

**Supplementary Fig. 5 | Expansion of quit metrics on Restaurant Row. a-d** Latency to quit in the wait zone from countdown onset (enter decision) to the moment of quitting and exiting the wait zone mid-countdown (a,c) across the entire Restaurant Row paradigm or (b,d) collapsed across the 1-30 s epoch split by flavor rankings (main effect of treatment: $F$=17.546, $p$<0.0001; VEH rank: $F$=10.541, $p$<0.0001; STZ rank: $F$=0.993, $p$=0.401). **e-f** Data from (b,d) split by the original starting offer preceding each quit event. **g-j** Data from (a-d) but instead plotting how much time left was remaining in the countdown at the moment of quitting. **k** Pooled data across all mice from the 1-30 s epoch showing the distribution of trial counts of quit events derived from each unique [time spent, time left] pairs across all permutations derived from trials of various starting offers, split by flavor rankings. **l** Pooled data across all mice from the 1-30 s epoch split by flavor ranking showing all quit events categorized in value terms, both the value of the original starting offer (offer value [Vo] = wait zone threshold – offer) and the value of the amount of time left remaining in the countdown at the moment of quitting (value left [$V_L$] = wait zone threshold – time left). Thus, this produces 4 quadrants, only three of which contain existing data. **m** Three categories of quit events based on offer value and value left terms. The green quadrant (Vo<0 & $V_L$<0) represent the most efficient quit decisions from economically disadvantageous offers that in theory should have been skipped instead. Efficient quits describe quitting fast enough before it otherwise would have been more consistent with crossing one's wait zone threshold to simply finish waiting instead. **n-o** Percentage of all quit events that derive from each quit category split by quit type in (n) or superimposed in (o) but split by groups of mice. Quit type x rank: $F$=49.958, $p$<0.0001. Note the interaction between quit type and treatment ($F$=8.645, $p$<0.001) but not schedule ($F$=0.044, $p$=0.957). These data do not capture the added change in quitting behavior that the sunk cost analysis captures (more dynamic changes due to the passage of time). Shaded / error bars represent ±1 SEM.

Supplementary Figure 6

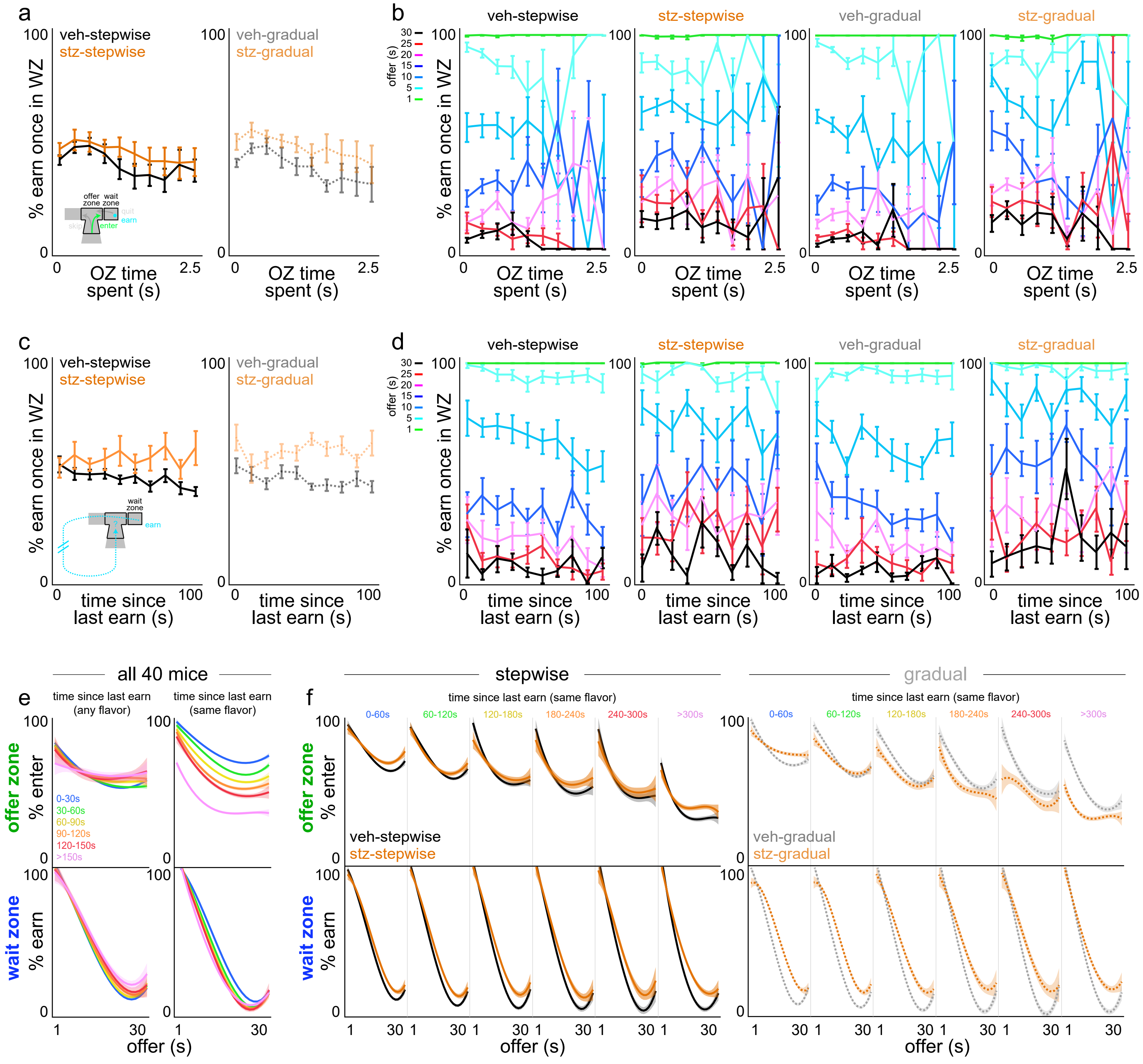

**Supplementary Fig. 6 | Valuing the passage of other forms of time on Restaurant Row. a-b** Time spent in the offer zone before making an enter decision (e.g., fast vs. slow decisions in the offer zone) has no significant bearing on the likelihood of staying vs. quitting in the wait zone once entered either (a) overall or (b) segmented by what the starting offer of the delay was in color (relatively flat horizontal lines for each color that scale vertically reflecting the overall likelihood of quitting low- vs. high-delay offers, $F$=0.571, $p$=0.450). Fewer samples exist at longer OZ time spent, as animals are more likely to skip in the offer zone instead. **c-d** Time spent navigating around the maze since last reward was earned of any flavor (e.g., recently ate vs. it has been some time) has no significant bearing on the likelihood of staying vs. quitting in the wait zone once entered either (a) overall or (b) segmented by what the starting offer of the delay was in color upon arriving in the wait zone. While there is an effect of treatment x time ($F$=10.457, $p$<0.01), this effect does not significantly present across offers ($F$=0.292, $p$=0.589) nor interact with schedule ($F$=1.627, $p$=0.202). **e-f** Interestingly however, we find more complex value structure in this analysis by (1) whether or not the behavior in question is offer zone enter decisions vs. wait zone earn decisions and (2) whether or not the time metric is time elapsed since last earn of any flavor vs. time elapsed since last earn of the same flavor (calculated here first split by flavor ranking but then re-collapsed across flavors to show more simplified summary metrics). Data here is also plotted against cued offer costs along the x-axis to show psychometric curves of offer zone enter decisions or wait zone earn decisions as a function of cost. In (e), across all 40 mice, we only observe modulation of choice behavior in the offer zone when time elapsed considers time spent since last earn of a specific flavor. In (f), splitting this across schedules, with separate curves plotted for each window of time elapsed since last earn of the same flavor, we observe modulation of offer zone choice behavior differently between VEH- and STZ-treated mice only in those tested on the gradual schedule. Error bars represent ±1 SEM. Shading represents 95% confidence interval of cubic fit.

Supplementary Figure 7

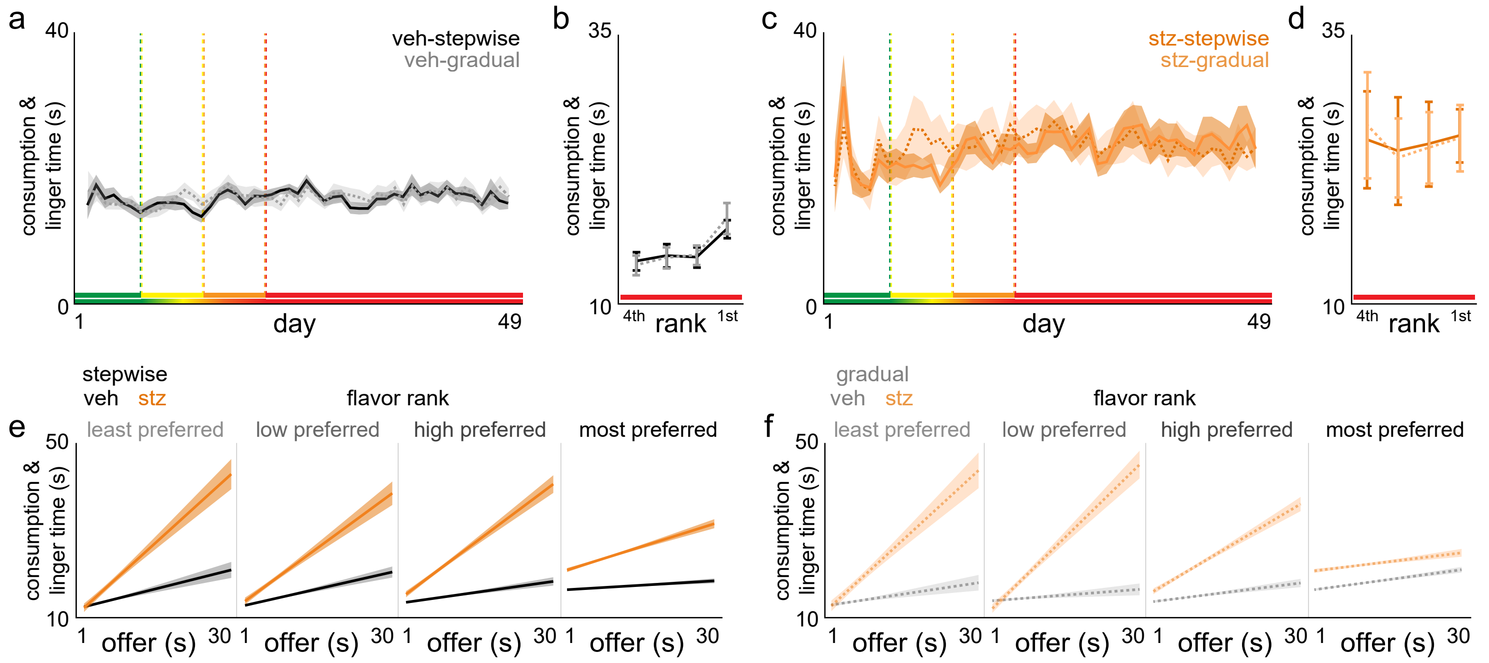

**Supplementary Fig. 7 | Post-earn behavior on Restaurant Row.** After mice earn a reward, time was measured from pellet delivery onset until mice left the feeder site and exited the wait zone. This encompasses time spent consuming the reward as well as, as previously reported, additional time mice spent lingering at the reward site without engaging in any overt feeding behavior. **a-d** Consumption and linger time plotted (a,c) across the entire Restaurant Row paradigm or (b,d) split by flavor rankings collapsed across the 1-30 s epoch. **e-f** Data in (b,d) split by the original starting offer of the cued cost required to earn the reward on each trial, split by flavor rankings. We previously reported this behavior includes a component of a within-trial conditioned-place-preference-like behavior that carries some hedonic value or appraisal of the reward just consumed, as well as is sensitive to the amount time invested prior to earning the reward, even within flavor. Shaded / error bars represent ±1 SEM. Shading in (e-f) represents 95% confidence interval of linear fit.

Supplementary Figure 8

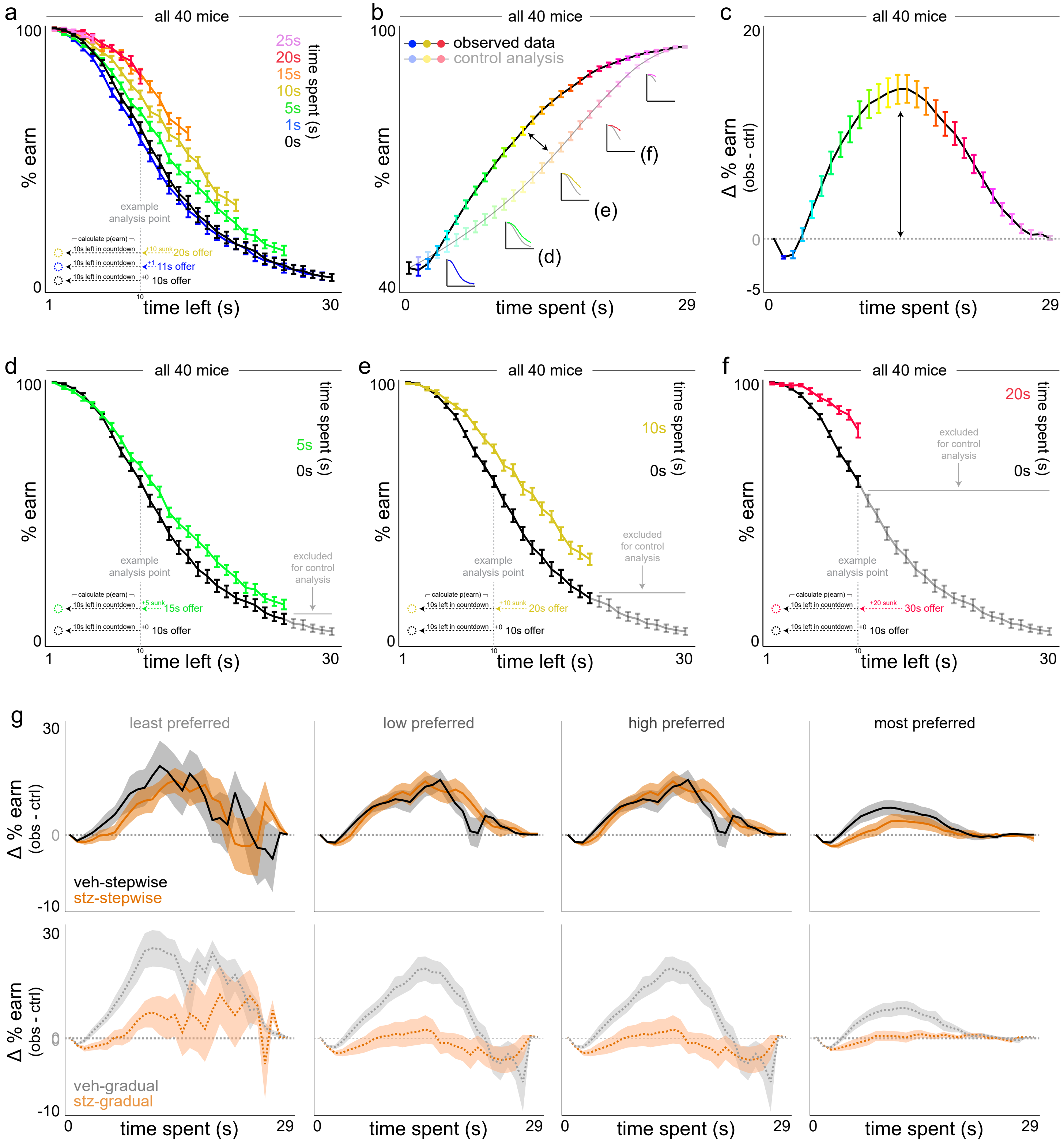

**Supplementary Fig. 8 | Visual explanation of sensitivity to sunk costs analysis in the wait zone and expanded data by flavor. a-c** Sunk cost analysis of staying behavior in the wait zone, demonstrated using all mice. (a) The likelihood of staying in the wait zone and earning a reward (e.g., not quit) is plotted as a function of time left in the countdown along the x-axis and time already spent waiting orthogonally in color. Note the black 0 s time spent curve represents animals having just entered the wait zone from the offer zone. Inset vertical dashed gray line illustrates an example analysis point comparing three sunk cost conditions originating from different starting offers but matched at 10 s left. Data from (a) dimensioned reduced in (b) collapsing across time left, instead highlighting the grand mean of each time spent sunk cost condition (color and x-axis). Insets depict data from curves in (a) are collapsed into the observed (sunk condition) and control (0 s condition) lines. Difference between curves in (b) are plotted in (c) in order to summarize the envelope of the overall effect of time already spent on escalating the commitment of staying in the wait zone. Horizontal dashed line represents 0. **d-f** Redisplay data from (a) but for three sunk cost conditions indicated in the insets in (b, 5 s green in d, 10 s gold in e, 20 s red in f) with the 0 sunk cost condition (black curve) repeated in each panel. Because each colored sunk cost curve derives from trials where the starting offer cost is higher than the matched time left value of the 0 s cost condition, some data points do not exist and are missing from the sunk cost curve on the rightward end. Thus, when reducing dimensions, to control for the inflated summary calculations of % earn due to missing data alone, the control analysis iteratively excludes matched data from the black 0 s sunk curve (highlighted in gray) before collapsing data to yield the resultant control curve in (b). **g** Delta curves for VEH- and STZ-treated mice split by schedule and flavor rankings. Shaded / error bars represent ±1 SEM.

### Supplementary Figure 9

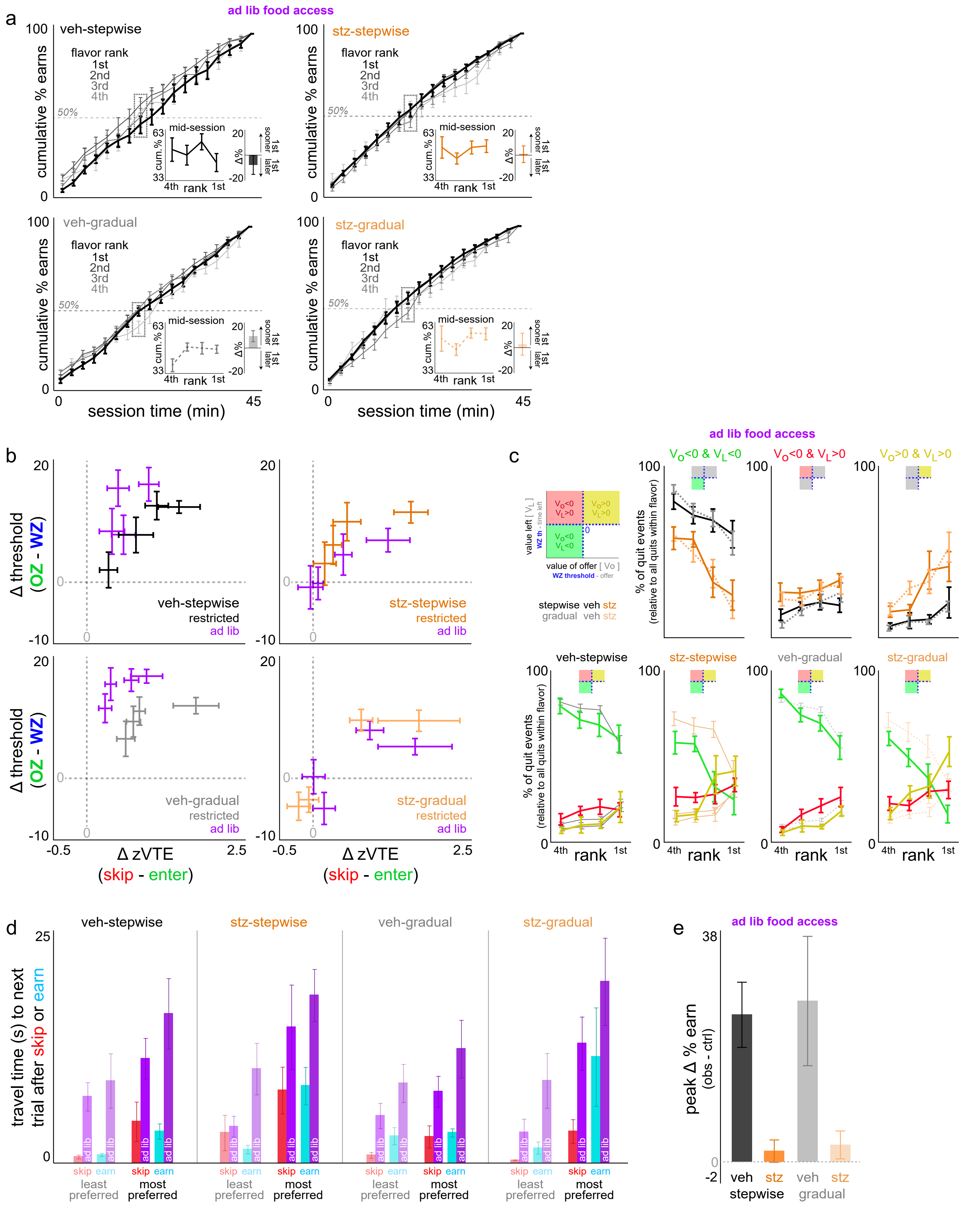

**Supplementary Fig. 9 | Expansion of additional Restaurant Row behavior while ad lib access to food was restored in the home cage. a** Meal consumption patterns as in Fig. 4e. Note the overall gross collapsing of flavor patterns across all groups (sign test mid-session delta score against zero: VEH-stepwise: $t$=-1.020, $p$=0.335; VEH-gradual: $t$=2.140, $p$=0.061; STZ-stepwise: $t$=0.721, $p$=0.490; STZ-gradual: $t$=0.203, $p$=0.843) compared to restricted patterns as in Fig. 4e. **b** Difference scores in vicarious trial and error (VTE) choice conflict in the offer zone skip minus enter plotted against difference scores in offer zone minus wait zone threshold policies as in Supplementary Fig. 4f, comparing food restricted baselines (color of each group) to ad lib data (purple). Flavor rankings split and superimposed on top of one another as in Supplementary Fig. 4f (no flavor-specific visual indicators are labeled intentionally to highlight the gestalt of bidirectional shifts among VEH- vs. STZ-treated mice driven largely by changes only in wait zone thresholds and not offer zone thresholds (largely a y-axis shift), with relatively less changing along the x-axis in VTE (although with some decrease observable in VEH-treated mice), as depicted in Fig. 8d-f. **c** Categorization of quit types as in Supplementary Fig. 5m-o. Top shows data split by quit type, only depicting ad lib access data only. Bottom superimposes quit type separated by groups of mice. For visual reference, superimposed in the background in thinner lines in the color of each group of mice is the baseline restricted data while the ad lib data is plotted in the color of each quit type (green, gold, red) to appreciate no changes in VEH-treated mice while there are changes in STZ-treated mice as a result of ad lib access. Of note, STZ-gradual mice show the greatest change in distribution of quit types particularly for most preferred flavors. **d** Differences in travel time between restaurants in the hallways split by groups of mice, after exiting least (faint) vs. most (opaque) preferred restaurants, and immediately after just skipping (red) or earning and consuming a reward (cyan). Restricted baseline data in red/cyan while the corresponding ad lib data is adjacent in purple. Note the overall slowing in travel times due to ad lib access ($F$=22.657, $p$<0.0001). Also note the value structure in anticipatory travel times that interact with heading toward the next flavor after leaving least (faster) vs. most preferred restaurants (slower, $F$=49.402, $p$<0.0001), in spite of global differences in speed after skipping (faster) vs. having just eaten (slower, $F$=5.984, $p$<0.05). **e** Larger visual redisplay of the same data plotted as insets in Fig. 8g. Error bars represent ±1 SEM.

Supplementary Figure 10

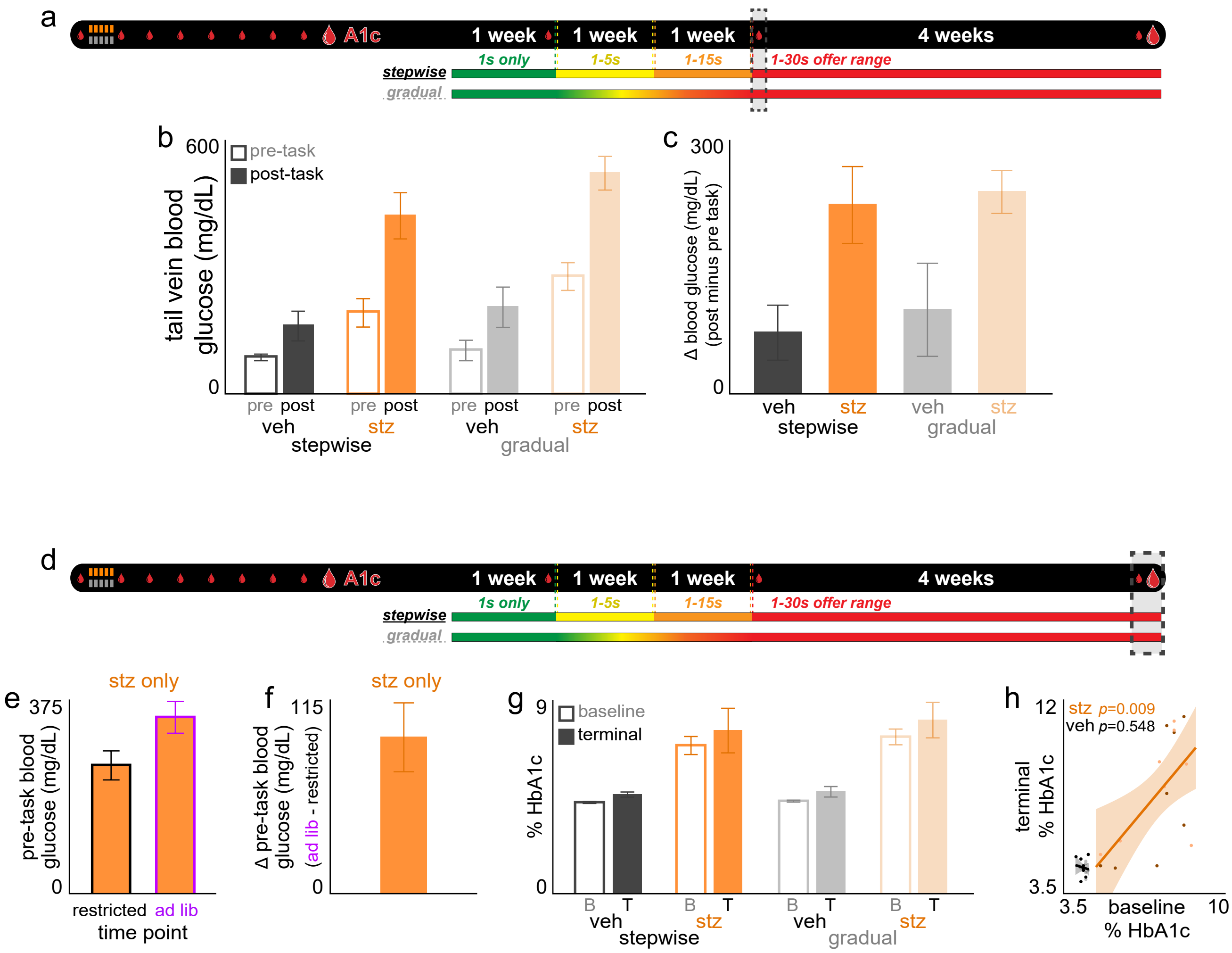

**Supplementary Fig. 10 | Additional timepoints of sampling blood glucose levels longitudinally across the entire Restaurant Row paradigm. a** Timeline. Dashed box indicates time period (day 22) relevant for panels (b-c). This includes tail vein pre- and post-task samples of blood glucose levels (small droplet icon) for all 40 mice. **b** Pre- and post- task blood glucose levels. **c** Change in blood glucose levels post minus pre task. **d** Timeline. Dashed box indicates time period relevant for panels (e-h). This includes the final day of the experiment before sacrificing mice to extract terminal HbA1c levels (large droplet icon) as well as additional samples of tail vein pre-task only blood glucose levels obtained from STZ-mice only during the additional days of testing with ad lib access to regular chow after day 49 of the main experiment. **e** Pre-task blood glucose levels comparing ad lib access to restricted [using day 22] only in STZ-mice sampled at both timepoints. **f** Change in blood glucose from (e) ad lib minus restricted [pre-task only]. **g** HbA1c levels measured at baseline (before ever starting the entire Restaurant Row paradigm but after the initial 8-week STZ incubation [first large droplet in the timeline]) and terminally [second large droplet in the timeline]. **h** Data in (g) replotting time points against one another for all mice (darker dots visualize stepwise schedule). Error bars represent ±1 SEM. Shading represents 95% confidence interval of linear fit.
